# An ER Retrieval-and-Release System Reveals Ligand-Independent Basal Signaling by Cell Surface TCRs

**DOI:** 10.64898/2026.10.13.758962

**Authors:** Claudia Juraske, Simon Beyer, Sara Hartmann, Daniel Peschkov, Emmanuelle Moraes, Hamidreza Ebrahimiyan, Katharina Ruoff, Felix Rainer Schmidt, Anna Morath, Manuel T. Dorer, Friedel Drepper, Balbino Alarcon, Gina Fiala, Mohsen Basiri, Björn Lillemeier, Pitter F. Huesgen, Viviane Timmermann, Wolfgang W. Schamel

## Abstract

The T cell receptor (TCR) machinery comprises the ligand-binding TCRαβ and the CD3 signaling dimers. Individual subunits are synthesized in the endoplasmic reticulum (ER) and only the fully assembled TCRs reach the cell surface. We engineered a system to retained CD3ε is in the ER. This allowed only partial TCR assembly, while continuous retrieval of CD3ε from the Golgi back to the ER enabled complete assembly. Synchronous TCR release by biotin addition caused a transient increase in surface TCR abundance that gradually returned to steady-state levels. Mathematical modeling revealed that TCR levels were controlled by feedback mechanisms. Notably, surface-localized, but not ER-localized, TCRs spontaneously generated ligand-independent basal signaling. Phosphoproteomic analyses revealed that this signal is similar, but not identical, to ligand-induced TCR activation signals, with distinct phosphorylation patterns already evident at the CD3 subunits. Our findings establish the TCR retrieval-and-release system as a tool to investigate ligand-independent basal TCR signaling.

## Introduction

The T cell receptor (TCR) is a complex macromolecular machinery consisting of the transmembrane TCRαβ, CD3 εδ, CD3εγ and CD3ζζ dimers and is responsible for the activation of T cells in response to antigen binding to TCRαβ ^1,2^. The TCR has a stoichiometry of αβɛγɛδζζ ^3^ and only completely assembled TCRs are transported to the cell surface ^4–6^.

The individual subunits are co-translationally translocated into the ER and fold while forming the TCRαβ, CD3εδ, CD3εγ and CD3ζζ dimers ^7^. Then, CD3εγ and CD3εδ associate with TCRαβ into a partial αβεδεγ complex ^8–10^, which then associates with CD3ζζ to form the complete αβεδεγζζ TCR. Where this last assembly step occurs is still unresolved. When CD3ζζ is absent, the partial TCR complex is transported to the Golgi, and CD3ζζ likewise reaches the Golgi when other subunits are missing, suggesting that the final assembly step could occur there ^11^. However, because all subunits are synthesized in the ER, CD3ζζ may instead associate with the partial complex in the ER. Supporting this, complete TCR assembly was observed in B cell-derived ER vesicles *in vitro* ^12^.

Antigen binding to TCRαβ leads to the phosphorylation of the CD3 cytoplasmic tails, initiating downstream signaling and T cell activation ^13,14^. The CD3ɛ, CD3γ, and CD3δ tails contain each one immunoreceptor tyrosine-based activation motif (ITAM), whose characteristics are two spaced tyrosines, while CD3ζ contains three ITAMs. Phosphorylation of these tryosines is accomplished by the Src family kinase LCK and double phosphorylation leads to the recruitment of the kinase ZAP-70 ^15^. Downstream of ZAP-70 are adaptor proteins, such as LAT, and classical signaling molecules, such as PLCγ1 and the MAP kinases ERK.

Several models on the molecular mechanisms of how antigen binding (or anti-CD3 antibody binding) causes CD3 ITAM phosphorylation have been proposed ^14,16,17^. In the allosteric model the activity of the TCR is regulated by CD3 conformational changes ^18–21^. According to this model the individual TCRs exist either in the resting, inactive or in the active conformation. In the resting conformation the CD3 cytoplasmic tails are not exposed and thereby the signaling motifs, such as the ITAMs, are not accessible. Upon antigen or antibody binding, the active conformation of the TCR is stabilized and the CD3 motifs are exposed allowing LCK to phosphorylate and signaling molecules to bind ^19,22^.

Two classes of antigen have been identified: i) those with high affinity for the TCR. They strongly activate the signaling pathways and their associated transcription factors, ultimately promoting T cell activation and initiating an immune response. ii) those with low affinity for the TCR. They induce tonic signaling, which is characterized by low-level phosphorylation of CD3 ^23^ and weak signaling, which does not cause T cell activation ^24^. Tonic signals contribute to T cell homeostasis ^25^ and impact the sensitivity of the TCR for high-affinity antigens ^26,27^. T cells distinguish between high-affinity and low-affinity ligands by kinetic proofreading at the TCR ^28,29^.

Besides these ligand-induced signaling, ligand-independent basal signaling exists in T cells ^30,31^. This signal serves to repress the transcription of the RAG genes and promotes the transcription of TCRα. Although this signal uses the TCR-downstream signaling machinery, such as LAT, it was argued that it occurs independent of the TCR, since Jurkat T cells without a fully assembled TCR on the cell surface also show this basal signal ^30^. However, partial assembled TCR complexes could have been involved in generating the basal signal (Swamy *et al*, 2016).

To investigate TCR assembly and the generation of basal signaling, we engineered two systems in which TCRs are retained inside the cell and released for transport to the cell surface upon biotin addition. These systems, referred to as the TCR retention/retrieval-and-release systems, enabled us to control TCR localization and investigate whether TCRs can initiate basal signaling. We found that basal signals originate at the TCR, but only if the TCR is localized at the cell surface and not at the ER membrane.

## Results

### In the Ii-εRUSH ER retention system, εRUSH is retained inside the cell, but TCRs hardly reach the cell surface upon biotin addition

To experimentally modulate TCR surface levels, we sought to establish a system with which we can retain TCRs in the ER and release them for transport to the cell surface on demand. We made use of the published RUSH system ^32^. In our case, human CD3ε was fused at its ectodomain to SBP and moxGFP ^33^, referred to GFP here, resulting in the SBP-GFP-CD3ε construct, called εRUSH (Fig. 1A). To retain εRUSH in the ER, we used an ER-resident protein – the human invariant chain of the major histocompatibility complex class II (Ii) – and fused it to streptavidin (Str) and the HA tag, resulting in the Ii-Str-HA construct ^32^, further called Ii-hook (Fig. 1A).

**Figure 1.**
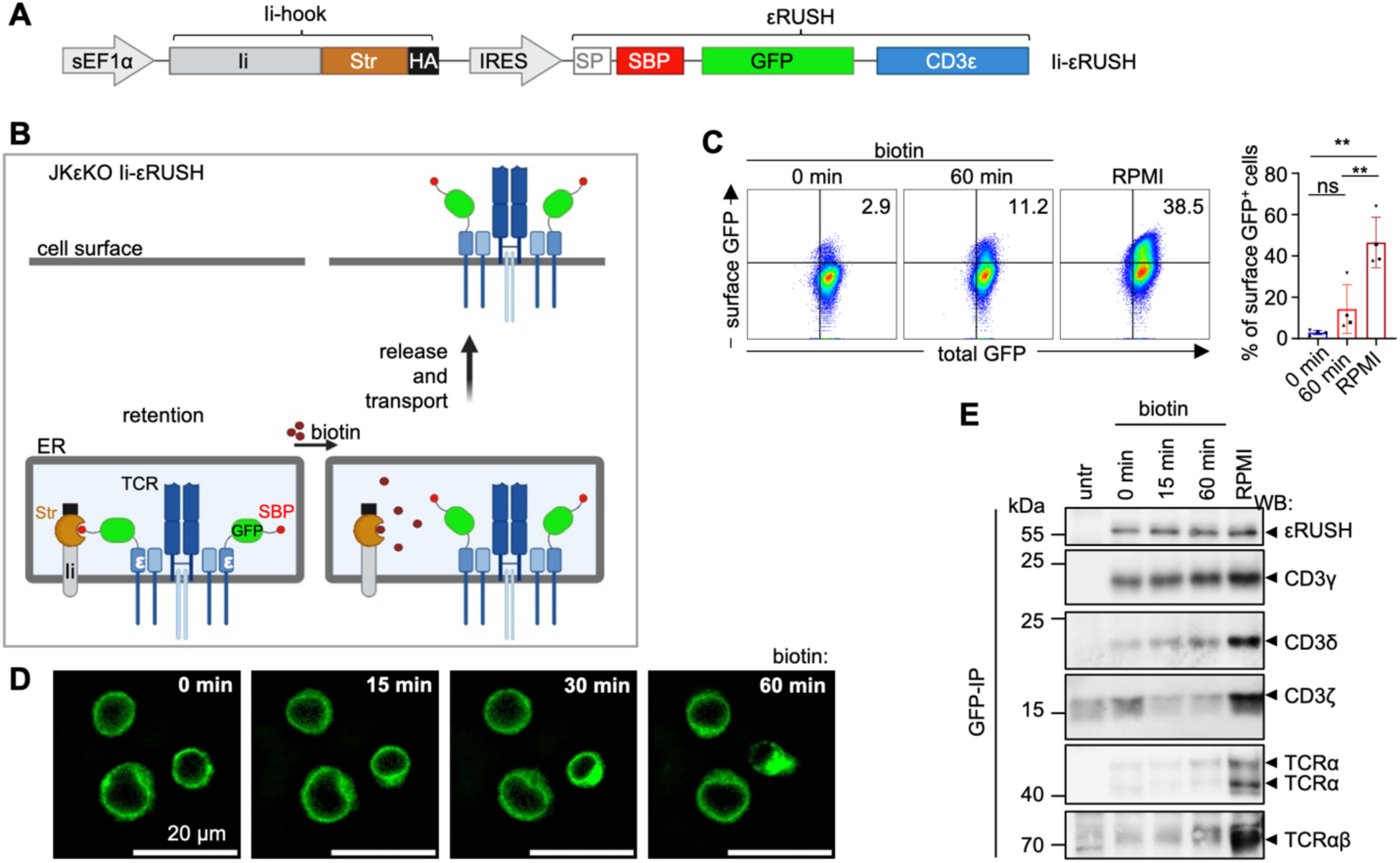
In the Ii-εRUSH system TCRs do not efficiently reach the cell surface after biotin-mediated release. **(A)** Scheme of the DNA encoding for the Ii-εRUSH construct. sEF1α: short EF1α promoter; Ii: MHCII invariant chain; Str: streptavidin; HA: HA tag; IRES: internal ribosomal entry site; SBP: Str-binding peptide. **(B)** Idea of the Ii-εRUSH system in JKεKO cells. **(C)** JKεKO Ii-εRUSH cells were cultured without biotin and left untreated (0 min), treated for 60 min with 40 µM biotin or cultured in biotin-containing RPMI medium (RPMI). Cells were stained with anti-GFP and secondary AF680-labeled antibodies and TCR surface levels were measured by flow cytometry. Percentages of surface GFP+ cells are plotted. Significance was analyzed by RM one-way ANOVA (n=4; 3 different transductions). **(D)** JKεKO Ii-εRUSH cells were cultured in medium without biotin and treated with 40 µM biotin while recording the GFP fluorescence by life microscopy. Pictures after the indicated times of biotin addition are shown (n=3). **(E)** JKεKO Ii-εRUSH cells were cultured in medium without biotin and left untreated (w/o) or treated for different times with 40 µM biotin. Untransduced JKεKO cells (untr) serve as a control. Cells were lysed and proteins purified by an anti-GFP immunoprecipitation (IP). Purified proteins were separated by reducing (top 5 panels) and non-reducing (lowest panel) SDS-PAGE. WB was done using antibodies directed against GFP (εRUSH), CD3γ, CD3δ, CD3ζ and TCRα (n=3).

SBP binds streptavidin, and thus, εRUSH will be retained in the ER. We expected that in the presence of the other TCR subunits the complete TCR would assemble and be retained in the ER (Fig. 1B). Upon addition of biotin, εRUSH and εRUSH-containing TCRs should be released from the Ii-hook and we anticipated that the TCRs would leave the ER being transported to the plasma membrane (Fig. 1B). The ER retention-and-release system for the TCR is further called the Ii-εRUSH system.

We designed a lentiviral Ii-εRUSH construct encoding the Ii-hook and εRUSH separated by an internal ribosome entry site (IRES, Fig. 1A). This construct was lentivirally transduced into Jurkat T cells in which the gene of CD3ε was deleted ^34^, called JkεKO Ii-εRUSH cells.

Cells were cultured in biotin-free medium, and biotin was added for 0 or 60 min (Fig. 1C). As control, cells were cultured in biotin-containing RPMI medium, further referred to as RPMI. All cells were GFP-positive, indicating that all cells were transduced and expressed εRUSH (Fig. 1C). TCR expression on the surface was measured by staining with an anti-GFP antibody that binds to εRUSH. Without biotin TCRs could be detected on the surface in 2.9% of the cells (0 min biotin treatment, Fig. 1C), indicating that εRUSH was retained inside most cells. With biotin only 11% (60 min biotin) of the cells had TCRs on their surface, indicating that release and transport to the cell surface was very inefficient. Similar results were obtained when the TCR on the surface was stained with anti-CD3ε or anti-TCRβ antibodies (Fig. S1A). The percentage of TCR surface expression varied between different transductions but was always very low in the presence of biotin. Under constant biotin presence (RPMI) many more cells expressed a TCR on their surface (39%, Fig. 1A and S1A).

As controls, we show that the Ii-hook is expressed and that the expression of both the Ii-hook and the total εRUSH amount (total GFP) stayed the same without and with biotin treatment (Fig. S1B). Next, we followed the GFP signal of εRUSH via fluorescence microscopy in living JkεKO Ii-εRUSH cells upon addition of biotin (Fig. 1D and movie 1). In nontreated cells, εRUSH and εRUSH-containing complexes were localized to structures most likely being the ER. After addition of biotin, the localization of εRUSH did not change in most cells, being in line with our finding that TCR do not appear on the cell surface. In those few cells where TCRs did reach the surface (Fig. 1C), the amount might have been too small to be seen in microscopy.

We thought that the two εRUSH proteins per TCR might hinder TCR assembly and/or transport to the cell surface. Therefore, we also tested the Ii-εRUSH system in JK wild-type (WT) cells expressing endogenous CD3ε, in which some of the TCRs should contain one εRUSH and one CD3ε WT protein. To only detect εRUSH-containing TCRs, we stained with an anti-GFP antibody. However, in the presence of biotin even fewer cells expressed the TCR on the cell surface compared to JkεKO Ii-εRUSH cells (Fig. S1C).

Increasing the concentration of biotin did not lead to increased expression of the TCR on the surface of JkεKO Ii-εRUSH cells (Fig. S1D), indicating that our concentration of biotin (40 µM) was not limiting. Indeed, this concentration was used in the original publication on the RUSH system ^32^.

In conclusion, with the Ii-εRUSH system, the εRUSH construct is retained well inside the cells, but does not efficiently come to the surface with biotin.

### During ER retention using the Ii-εRUSH system, TCRs assemble incompletely

After biotin addition, most εRUSH remained in the ER (Fig. 1D and movie 1). These might be εRUSH molecules that are not assembled to a complete TCR competent for export. We assumed that TCRs would completely assemble to the αβεδεγζζ complex during ER retention (Fig. 1B), however, this might not have been the case. To test this, JkεKO Ii-εRUSH cells were treated without (0 min) and with biotin for different times (15 and 60 min) and an anti-GFP IP was performed to purify εRUSH-containing complexes (Fig. 1E). As a negative control, untransduced JkεKO cells were used (untr). During retention in the ER, when cells are in medium without biotin, we could detect εRUSH and CD3γ, indicating that εRUSH-CD3γ dimers formed. CD3δ was only co-purified in small amounts, indicating that εRUSH-CD3δ dimers formed inefficiently. Likewise, TCRα and TCRβ hardly assembled with εRUSH (TCRβ was detected via its heterodimerization with TCRα, last panel). The presence of a background signal in untransduced cells prevents definitive conclusions about CD3ζ. Anyhow, the complete TCR did not assemble and only partial complexes have been formed when εRUSH was retained in the ER.

When the JkεKO Ii-εRUSH cells were continuously cultured in medium containing biotin (RPMI), so that εRUSH was never retained in the ER larger amounts of all subunits, including CD3δ, TCRα, TCRβ and CD3ζ could be co-purified with εRUSH, suggesting that some complete TCRs formed (Fig. 1E, RPMI). Indeed, in this case many more TCRs could be detected at the cell surface (Fig. 1C) that contained CD3 dimers and the TCRαβ dimer (Fig. S1A). Hence, it might not have been the fusion of SBP and GFP to CD3ε, but a lack to travelling to the Golgi, that prevented assembly of the TCR. Indeed, with a fusion of other large domains to CD3ε, such as a single chain Fv, the TCR assembled and was transported to the cell surface ^34–36^.

In conclusion, in the Ii-εRUSH system without biotin, εRUSH is retained in the ER. During retention only partial TCR complexes assemble and hence after addition of biotin TCRs are hardly transported to the cell surface. Therefore, it might be that complete TCR assembly requires that the subunits and/or partial complexes leave the ER.

### In the KDEL-εRUSH ER retrieval system, εRUSH is retained inside the cell and TCRs are released to the cell surface upon biotin addition

In our ER retention system using the Ii-hook, the TCRs were not efficiently released to the cell surface, most likely due to partial TCR assembly. Next, we tested a system in which TCRs are retained in the cells by a retrieval mechanism. In this approach, we used the KDEL retrieval signal ^37^ fused to streptavidin, resulting in the Str-KDEL fusion construct ^32^, further called the KDEL-hook (Fig. 2A). The εRUSH was the same as above. With this KDEL-εRUSH system, we expect TCRs to escape from the ER and be retrieved back to the ER from the cis-Golgi preventing surface expression (Fig. 2B). Upon biotin addition TCRs should be released from the KDEL-hook and transported to the plasma membrane.

**Figure 2.**
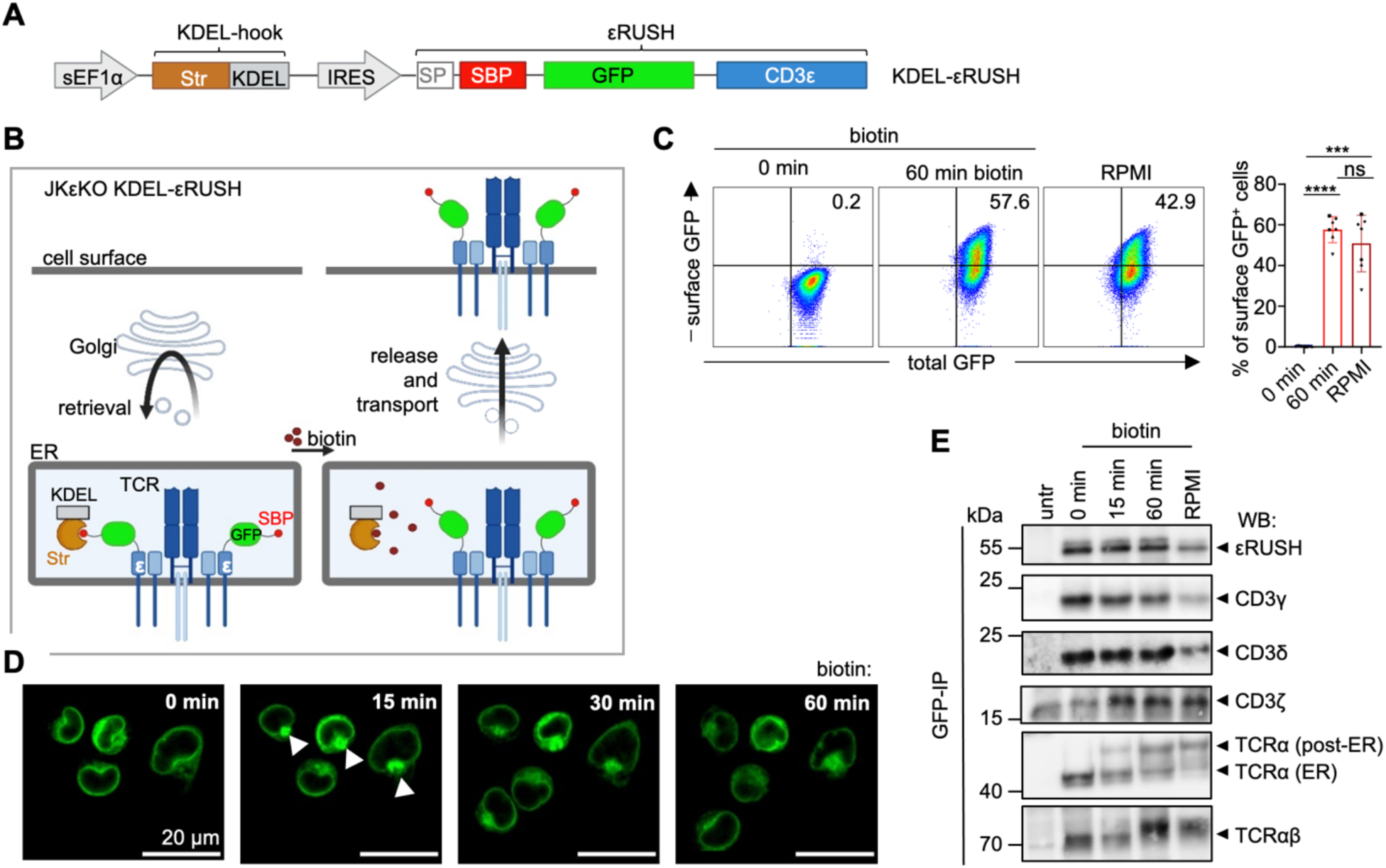
In the KDEL-εRUSH system TCRs reach the cell surface after biotin-mediated release. **(A)** Scheme of the cDNA encoding for the KDEL-εRUSH construct. KDEL: ER retrieval signal. **(B)** Scheme of the KDEL-εRUSH system in JKεKO cells. **(C)** JKεKO cells lentivirally transduced with the KDEL-εRUSH construct were cultured without biotin and left untreated (0 min) or treated for 60 min with 40 µM biotin or cultured in biotin-containing RPMI medium (RPMI). Cells were stained with anti-GFP and secondary AF680-labeled antibodies and TCR surface levels were measured by flow cytometry. Percentages of total GFP+ surface GFP+ cells are plotted. Significance was analyzed by ordinary one-way ANOVA (n=3; one transduction). **(D)** JKεKO KDEL-εRUSH cells were cultured in medium without biotin and treated with 40 µM biotin while recording the GFP fluorescence by life microscopy. Pictures after the indicated times of biotin addition are shown (n=3). **(E)** JKεKO KDEL-εRUSH cells were cultured in medium without biotin and left untreated (0 min) or treated for different times with 40 µM biotin. Untransduced JKεKO cells (untr) serve as a control. Cells were lysed and proteins purified by an anti-GFP IP. Purified proteins were separated by reducing (top 5 panels) and non-reducing (lowest panel) SDS-PAGE. WB was done using antibodies directed against GFP (εRUSH), CD3γ, CD3δ, CD3ζ and TCRα (n=3).

To test this new approach, JkεKO cells were lentivirally transduced with the KDEL-εRUSH construct, called JkεKO KDEL-εRUSH cells. TCR surface expression was measured by detecting εRUSH-containing TCRs with an anti-GFP antibody in cells without biotin (0 min) and after addition of biotin for 60 min, as well as in cells cultured in RPMI medium (Fig. 2C). TCRs were retained inside the cell without biotin and released to the cell surface in 58% of the cells after addition of biotin for 60 min. Fewer cells (43%) expressed the TCR on their surface in RPMI medium. The same trend was also seen by anti-CD3ε (Fig. S2A) and anti-TCRβ stains (Fig. S2A), suggesting was it was the assembled TCR that reached the surface and not εRUSH alone. Thus, with the KDEL-εRUSH system, the εRUSH construct is retained inside the cell, and reaches the surface with biotin treatment.

The surface expression of the εRUSH molecule was decreased after 60 min of biotin treatment compared to untreated cells (Fig. S2B). We will come back to this finding in the next figure.

We also tested the KDEL-εRUSH system in JK WT cells. Although surface TCR expression in the presence of biotin was similar to that in JkεKO KDEL-εRUSH cells, intracellular retention in the absence of biotin was less efficient (Fig. S2C), most likely due to endogenous CD3ε. Thus, we decided to use the JkεKO KDEL-εRUSH cells for further studies with the KDEL-εRUSH system. Finally, the GFP signal of εRUSH was monitored in living JkεKO KDEL-εRUSH cells using fluorescence microscopy after addition of biotin (Fig. 2D and movie 2). In untreated cells, εRUSH was localized to structures reminiscent of the ER. 15 min after addition of biotin, some GFP signal was visible in the Golgi complex (arrowheads) and some was still localized to the ER, as shown for the Ii-εRUSH system. The GFP signal in the Golgi complex decreased after 60 min after biotin addition. We could not detect TCR surface expression by fluorescence microscopy, since the plasma membrane cannot be distinguished well from the cytosol in T cells. However, TCR surface expression was shown by flow cytometry (Fig. 2C and S2A). Hence, we conclude that εRUSH was transported through the Golgi to the plasma membrane within 60 min. This kinetics was also seen for other proteins using the RUSH system ^32^.

In conclusion, with the KDEL-εRUSH system, εRUSH and εRUSH-containing TCRs were retained inside the cell in the absence of biotin. After addition of biotin, the TCRs were released in a synchronized manner, travelled through the Golgi and then reached the plasma membrane after 60 min. Some εRUSH molecules stayed in the ER after biotin addition; thus, not all εRUSH molecules might have assembled to complete TCRs in the ER.

### In the KDEL-εRUSH system, TCRs assemble during ER retrieval

With our retrieval-and-release system, TCRs were transported to the cell surface after biotin-mediated release (Fig. 2B, C and S2A). This might have been enabled by the full assembly of the TCR when retrieved from the cis-Golgi. To test this, εRUSH was purified from the JkεKO KDEL-εRUSH cells using an anti-GFP IP followed by SDS-PAGE and WB to probe for associated TCR subunits (Fig. 2E). During retrieval to the ER, when cells were cultured without biotin (0 min), we could detect the TCRα, TCRβ, CD3δ and CD3γ TCR subunits being co-purified with εRUSH. Due to the presence of a background band in the untransduced cells, no conclusions could be drawn regarding CD3ζ co-purification. However, when we purified εRUSH with an anti-CD3ε IP the CD3ζ subunit was present more when using the JkεKO KDEL-εRUSH cells than using the untransduced control (Fig. S2D), indicating that CD3ζ also assembled to the TCR. Upon release by addition, of biotin some TCRα molecules increased in size (Fig. 2E) being consistent with more complex glycosylation as obtained while traveling through the Golgi.

In conclusion, using the KDEL-εRUSH retrieval-and-release system, TCRs assemble into the complete αβεδεγζζ complex while being retained inside the cell. This allows them to be transported to the cell surface upon release from the KDEL-hook using biotin.

### Faster kinetics of TCR surface expression with the KDEL-εRUSH system compared to the Ii-εRUSH system

Having seen differences in TCR assembly and surface expression between the ER retention and the ER retrieval systems, we next investigated the kinetics of TCR surface expression in both systems. To this end, the JkεKO Ii-εRUSH and KDEL-εRUSH cells were cultured in medium without biotin, and biotin was added for different periods of time. We included cells that were always cultured in biotin-containing RPMI medium. TCRs on the surface were stained with an anti-GFP antibody that binds to εRUSH, and surface levels were quantified by flow cytometry (Fig. 3A and S3A). At most time points, the TCR surface levels of JkεKO Ii-εRUSH cells were lower than those of the JkεKO KDEL-εRUSH cells, being in line with full TCR assembly only in the KDEL-εRUSH system. Under continuous biotin exposure (RPMI), where εRUSH does not bind to any hook, TCRs in both systems should assemble and be transported to the surface similarly. Indeed, we found comparable levels of surface TCR expression in both systems (Fig. 3A).

**Figure 3.**
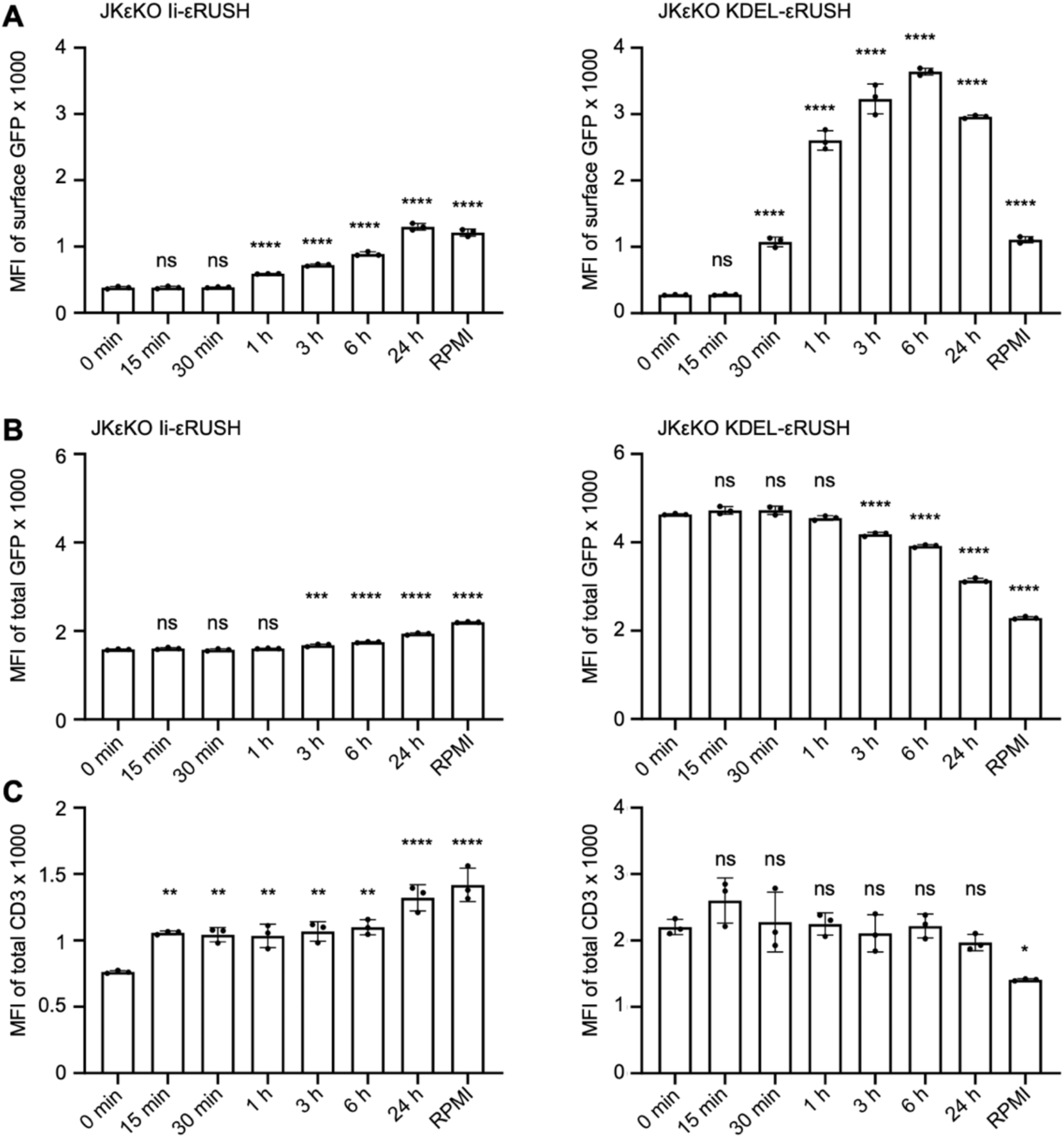
Kinetics of TCR surface expression in the Ii-εRUSH and KDEL-εRUSH systems. **(A)** JKεKO Ii-εRUSH (left) and JKεKO KDEL-εRUSH cells (right) were cultured in medium without biotin and left untreated (0 min), treated for different times with 40 µM biotin or were cultured in biotin-containing RPMI medium (RPMI). Cells were stained with an AF647-labeled anti-GFP antibody and TCR surface levels were measured by flow cytometry. The MFI of surface GFP is shown. Significance to the 0 min samples was analyzed by ordinary one-way ANOVA (n=3 for Ii-εRUSH, n>3 for KDEL-εRUSH; one representative experiment in triplicates is shown). **(B** and **C)** Cells as in (A) were fixed, permeabilized and stained with an BV421-labeled anti-CD3ε antibody and measured by flow cytometry. The MFI of total GFP is shown **(B)**, and the MFI of the anti-CD3ε stain is depicted **(C)**. Significance to the 0 min samples was analyzed by ordinary one-way ANOVA (n>3 for B and n=3 for C; one experiment in triplicates is shown).

In the Ii-εRUSH system, TCR surface levels slowly increased after biotin addition, being firstly detectable after 1 h and reaching the steady state at 24 h (Fig. 3A). This is consistent with our finding that during retention, complete TCRs cannot form, and suggests that after biotin addition, only newly synthesized TCRs can assemble and reach the surface. In contrast, in the JkεKO KDEL-εRUSH cells, the TCRs reached the surface already in 30 min after biotin-mediated release (Fig. 3A); thus, export-competent TCRs assemble during ER retrieval, leading to immediate transport to the cell surface upon biotin addition.

In JkεKO Ii-εRUSH cells, the total levels of εRUSH (total GFP) were lower during retention compared to the KDEL-εRUSH cells (Fig. 3B). This could be another sign of impaired assembly in the Ii-εRUSH system, since unassembled TCR subunits are degraded ^4,38^. With KDEL-εRUSH TCRs assembled and fewer individual subunits were degraded. Under constant biotin exposure (RPMI), where εRUSH does not bind to any hook, TCRs in both systems should assemble, and εRUSH degraded similarly. Indeed, we found comparable levels of total GFP in both systems (Fig. 3B, RPMI).

CD3ε pairs with CD3δ and CD3γ, and only then its extracellular immunoglobulin domain folds correctly ^39,40^ and can bind to the monoclonal anti-CD3ε antibody UCHT1 ^39,41^. In JkεKO Ii-εRUSH cells, correctly folded CD3ε (εRUSH) increased with time (Fig. 3C), suggesting that CD3ε that is retained in the ER by binding to Ii cannot efficiently pair with CD3δ and/or CD3γ. Once the CD3ε molecules are released, pairing increases. This is supported by the finding that in the presence of biotin εRUSH did not bind well to CD3δ (Fig. 1E) and that in the KDEL-εRUSH system, in which εRUSH is constantly retrieved from the Golgi in the absence of biotin, more folded CD3ε molecules were present (Fig. 3C) and εRUSH bound well to CD3δ and CD3γ (Fig. 2E).

When KDEL-εRUSH cells were treated with puromycin and new synthesis of TCRs was stopped before we added biotin, we did not see a difference in the amount of TCRs that reached the surface in 30 min compared to untreated cells (Fig. S3B). This shows that indeed the TCRs that were transported to the surface upon biotin addition within 30 min were those that were preformed inside the cell. After 60 min, only slightly more TCRs were detected at the cell surface without puromycin compared to the presence of puromycin, indicating that only a few TCRs were newly synthesized in one hour and reached the surface.

In the JkεKO KDEL-εRUSH cells, TCR surface levels increased up to 6 hours of biotin treatment and declined afterwards. This was seen with the anti-GFP and the anti-CD3ε stains (Fig. 3A and S3C). The increase is mostly due to surface transport of pre-assembled TCRs and only partially caused by newly synthesized TCRs (Fig. S4B). The following decrease in surface levels is in line with reduced total εRUSH levels over longer times (total GFP, Fig. 3B) and reduced folded εRUSH levels at longer time points (anti-CD3ε stain, Fig. 3C). In contrast, a decline in TCR surface or total εRUSH levels was not detected in the JkεKO Ii-εRUSH cells (Fig. 3A, B and C) and instead a slow constant increase is seen.

Together, this suggests that large amounts of completely assembled TCRs accumulate inside the cells – but only when TCRs are retained by retrieval from the Golgi back to the ER and not when they are retained in the ER. As a consequence of accumulating excess TCRs intracellularly in the KDEL-εRUSH system, too many TCRs reach the surface upon synchronous release; and with time the surface levels decline to a lower steady state. This is concomitant with the degradation of εRUSH after release, which might occur in lysosomes following spontaneous TCR internalization from the cell surface. From now on, we will call the JkεKO KDEL-εRUSH system the TCR retrieval-and-release system.

### T cells might regulate their TCR surface levels by control engineering

We sought to get insight into the mechanisms controlling TCR expression levels in our TCR retrieval-and-release system. To this end, we developed a mathematical model based on ordinary differential equations ^42^. The model describes TCRs in three compartments, the ER, cell surface and endosome. It further delineates the fluxes between these compartments along with the production in and degradation from the ER, export to the surface, internalization, recycling and degradation in lysosomes (Fig. 4A, solid lines), similar to earlier models ^43,44^.

**Figure 4.**
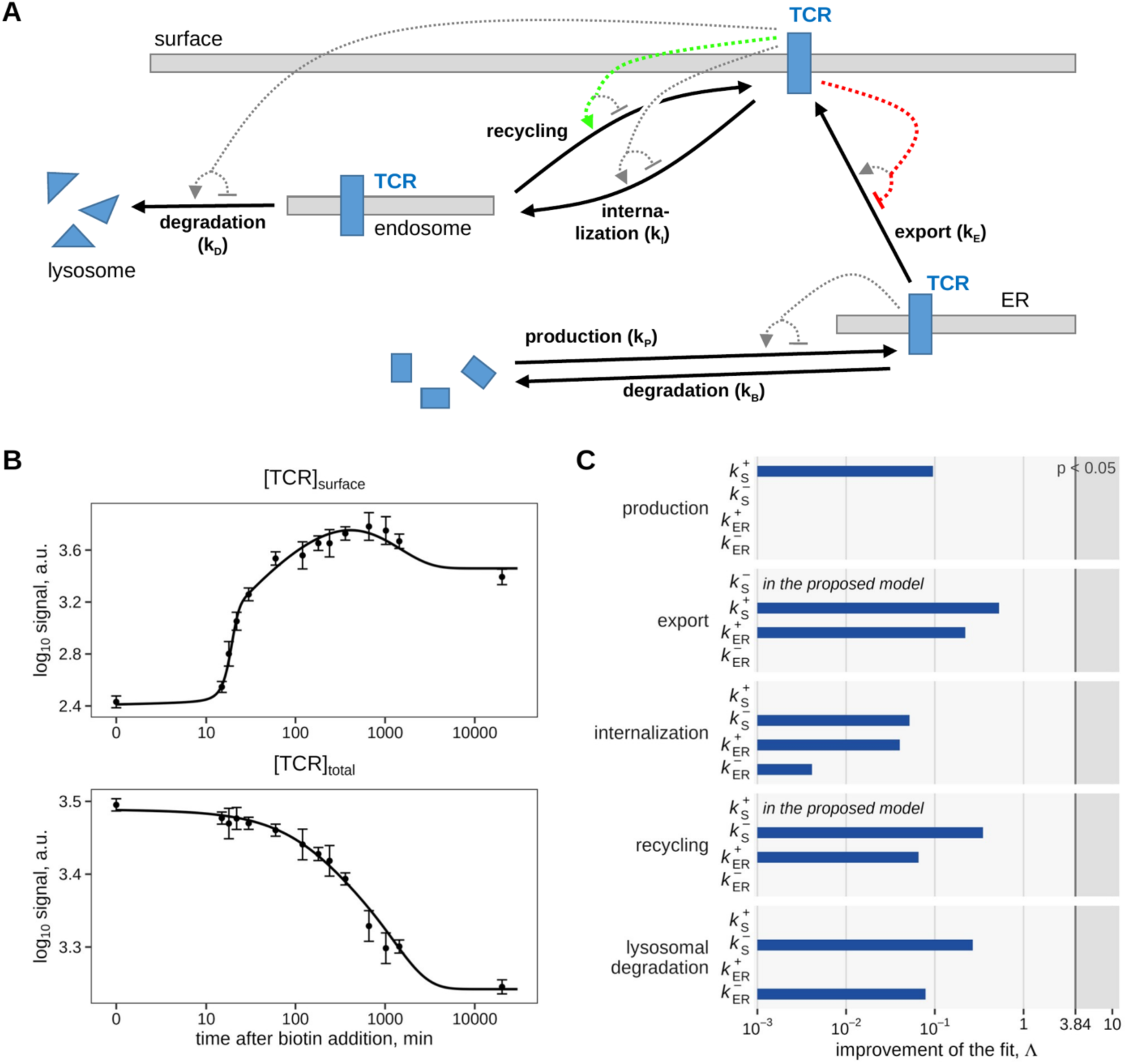
Mathematical model of TCR production, transport and degradation. **(A)** Scheme of the model. Solid arrows: fluxes; dotted arrows: feedbacks by surface and ER-localized TCRs that were tested (selection; all tested feedbacks are listed in C). The proposed model contains the inhibition of export by surface TCRs (red) and the promotion of recycling by surface TCRs (green). Recycling has no basal rate; it occurs only through its promotion by surface TCRs. Internalized TCRs that are not recycled are degraded after transport to lysosomes. **(B)** The proposed model (lines) and the flow cytometry data (mean ± s.e.) of surface TCR (top) and total TCR (bottom) after biotin-mediated release. The fit to the Western blot data are shown in the Supplement “Mathematical Modeling”. **(C)** Test of further feedbacks. Each promotion (k⁺) or inhibition (k⁻) of production, export, internalization, recycling or lysosomal degradation (arrow to the lysosome in (A) by surface (S) or ER-localized (ER) TCRs was added on its own to the proposed model. Bars: improvement of the fit by the added feedback (likelihood ratio statistic Δ). A feedback would be required if its bar reached the grey area (p < 0.05). Feedbacks without a visible bar improve the fit by less than 10⁻³. The two feedbacks of the proposed model have no bar, because they are already part of it. No further feedback improved the fit significantly.

The model was calibrated to the time courses of surface TCR and total TCR after biotin-mediated release, both measured via the GFP of εRUSH (Fig. 3), and to the amounts of ER-localized and post-ER TCRα (Fig. 2E). Post-ER TCRα migrates more slowly in SDS-PAGE than ER-localized TCRα, because its glycosylation gets more complex in the Golgi ^45,46^. Starting from the model without feedbacks (Fig. 4A, solid lines), we tested inhibitory and promoting feedbacks on TCR production, export, internalization, recycling and degradation by surface TCRs and ER-localized TCRs (Fig. 4A, dotted lines). A feedback term was included in the model only where justified by the data, based on likelihood ratio tests and profile likelihood evaluation ^47,48^. For details we refer to the Supplement “Mathematical Modeling”.

The data indicated that two feedbacks by surface TCRs are required: an inhibition of TCR export from the ER to the surface and a promotion of recycling, by which endosomal TCRs return to the surface. All parameters of this proposed model were identifiable, and removing either feedback resulted in a significantly worse fit (Supplement “Mathematical Modeling”).

Our model describes the time courses of both total TCR and surface TCR (Fig. 4B) as well as the amounts of ER-localized and post-ER TCRs (Supplement “Mathematical Modeling”). We then added each further feedback, by surface or ER-localized TCRs of any of the processes, to the proposed model one at a time. None of them improved the fit (Fig. 4C). In particular, the data did not require any feedbacks by ER-localized TCRs.

In conclusion, our modeling supports a key role of feedbacks by surface TCRs, but not by ER-localized TCRs, in maintaining surface receptor homeostasis.

### TCRs generate a basal signal after having left the ER

A feedback mechanism from surface TCRs would imply that the TCRs at the plasma membrane are signaling competent in the TCR retrieval-and-release system. To test this, we treated the JKεKO KDEL-εRUSH cells for 0, 1 or 16 hours with biotin, or grew them in biotin-containing RPMI medium. Then, TCRs were stimulated using anti-CD3ε antibodies for 6 additional hours. Upregulation of the activation marker CD69 was quantified by flow cytometry (Fig. 5A). Cells in which TCRs were not released to the surface (0 h biotin) did not upregulate CD69, being consistent with a lack of surface TCRs (Fig. 2C and S2A). In contrast, biotin-treated cells upregulated CD69 (Fig. 5A), showing that the surface TCRs were signaling competent.

**Figure 5.**
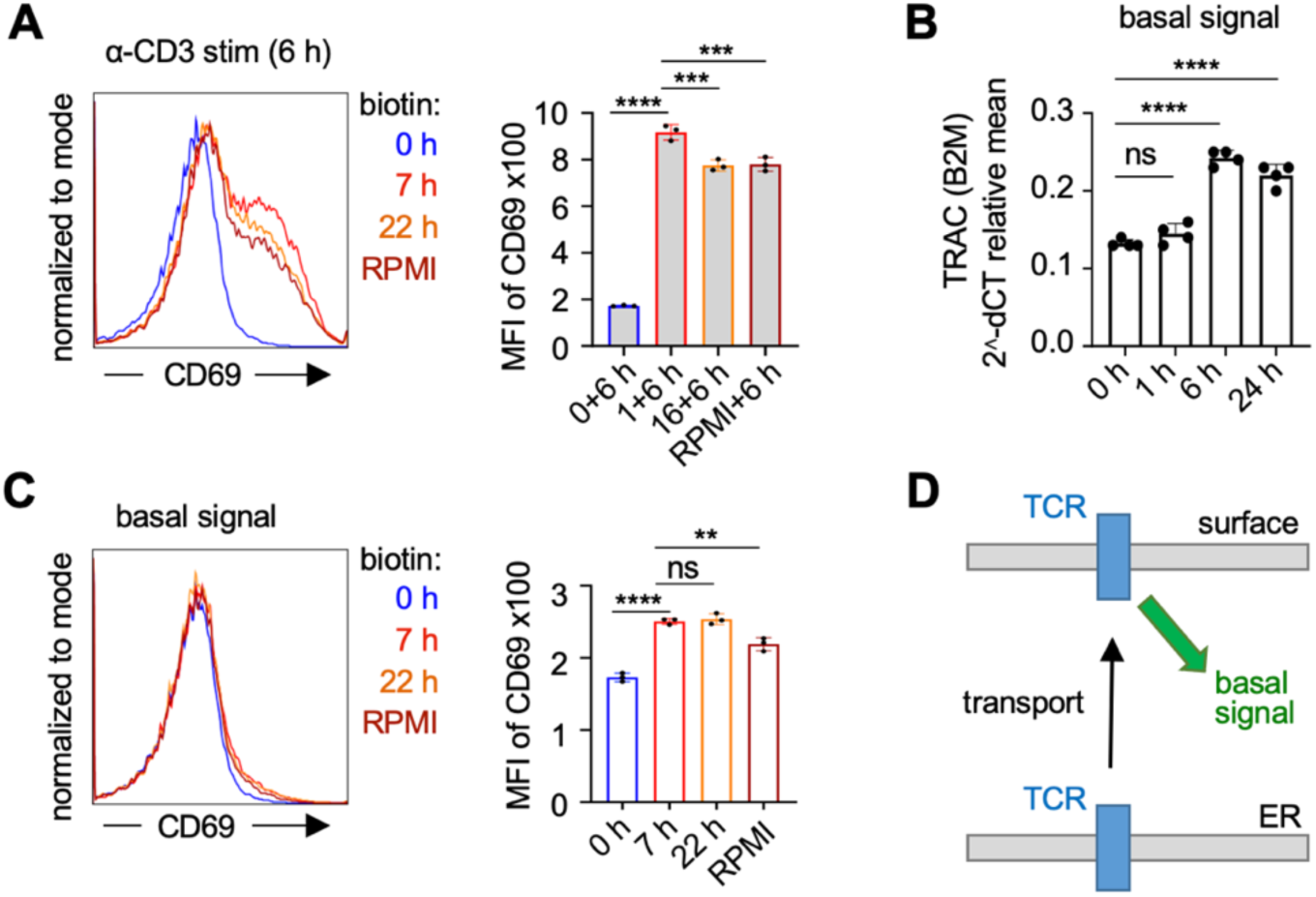
Antibody-induced and basal signaling of TCRs in the KDEL-εRUSH system before and after release. **(A)** JKεKO KDEL-εRUSH cells were cultured in medium without biotin (0 h) or treated for 1 or 16 hours with 40 µM biotin or were cultured in biotin-containing RPMI medium. Then cells were stimulated with 5 µg/ml anti-CD3ε antibody (α-CD3 stim.) at 37°C for 6 hours during which the biotin treatment was continued (except for the cells that were not treated with biotin). Cells were stained with an APC-labeled anti-CD69 antibody and measured by flow cytometry. Histograms and the MFI of CD69 are shown. Significance was analyzed by two-way ANOVA (n=2; shown one representative experiment in triplicates). **(B)** JKεKO KDEL-εRUSH cells were cultured in biotin-free medium and treated with 40 µM biotin for 1, 6, or 24 h, or left untreated. RT-qPCR was performed to quantify TCRα (TRAC) mRNA expression levels. The obtained Cq values were normalized to the housekeeping gene B2M. Each sample was analyzed in technical triplicates. Significance was analyzed by one-way ANOVA (n=3). **(C)** The experiment and biotin treatment was done as in (A), but leaving the cells unstimulated. Histograms and MFI of CD69 are shown. Significance was analyzed by two-way ANOVA (n=2; shown one representative experiment in triplicates). **(D)** Scheme showing that TCRs in the ER do not emit a basal signal, whereas TCRs at the cell surface do.

Importantly, a feedback from surface TCRs would also imply that these TCRs emit a signal in the absence of any ligand binding. We trapped the TCRs inside the cell using our TCR retrieval-and-release system, released them to the cell surface, and measured signs of the basal signal. One hallmark of ligand-independent signaling is the enhanced transcription of the mRNA encoding the TCRα subunit (TRAC) ^30,31^. Firstly, we performed real-time quantitative reverse transcription PCR (RT-qPCR) to measure TRAC mRNA levels using JKεKO KDEL-εRUSH cells treated or not with biotin. We found that TRAC mRNA levels were increased at 6 h of biotin treatment compared to 0 or 1 h (Fig. 5B). The same held true for cells cultured in RPMI. This correlated with the TCR levels on the cell surface (Fig. 3A), assuming that it takes a short while from the appearance of the basal signal to the accumulation of mRNA molecules to be detected.

It had been suggested that the basal signal might be similar to the ligand-induced signal, but weaker in its amplitude ^30,31^. Hence and secondly, we quantified CD69 expression using flow cytometry and JKεKO KDEL-εRUSH cells treated or not with biotin and left unstimulated (Fig. 5C). Compared to the cells without biotin treatment, cells with 7 and 22 hours of biotin treatment, in which TCRs have left the ER upregulated CD69 – although mildly, as expected from the basal signal. Cells constantly cultured in biotin-containing RPMI, in which fewer TCRs were on the surface than in the cells treated with biotin for 7 or 22 hours (Fig. 3A), showed less CD69 upregulation.

In conclusion, our data show that TCRs that have left the ER and presumably are on the cell surface generate a basal signal (Fig. 5D). This signal is not detectable with ER-localized TCRs.

### Enhanced total TCR presence upon ER retrieval in primary T cells

Using our TCR retrieval-and-release system, we found that the cells overproduce TCRs when TCRs were kept in the ER (Fig. 2 and 3). Next, we wanted to test whether this would also be the case in primary human T cells. To this end, we made use of the CD3ε-ER-retrieval (CD3ε-ERR) construct that we had developed earlier ^49^. In the CD3ε-ERR construct, the ER retrieval signal KKxx ^50^ is directly fused to the cytosolic tail of CD3ε (Fig. 6A). Hence, TCRs that incorporate this molecule would be localized to the ER by a retrieval mechanism from the cis-Golgi (Fig. 6B).

**Figure 6.**
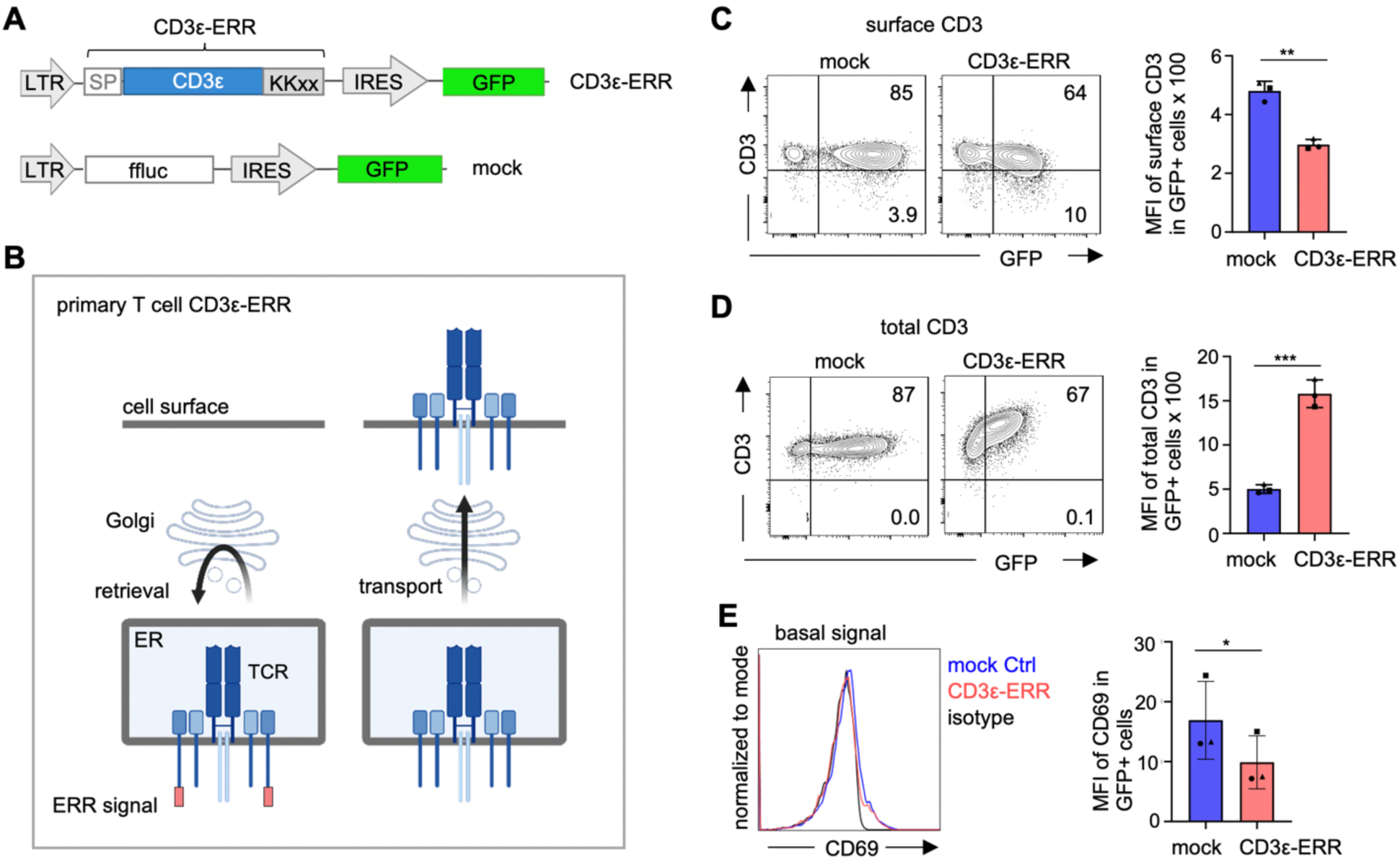
Expression of CD3ε-ERR shows that surface TCRs generate a basal signal in primary human T cells. **(A)** Scheme of the DNA encoding for the CD3ε-ERR and control constructs. LTR: long terminal repeat promoter; ERR: ER retrieval; ffluc: firefly luciferase. **(B)** Principle of the CD3ε-ERR system; please note that in the primary T cells wild type CD3ε and CD3ε-ERR are co-expressed. **(C)** Expanded human T cells from blood were retrovirally transduced to express mock or CD3ε-ERR. After five days of culture cells were stained with APC-labeled anti-CD3 antibodies and TCR surface levels were measured by flow cytometry. The MFI of CD3 in the GFP+ cells is shown. **(D)** Cells as in (C) were fixed, permeabilized and stained with an APC-labeled anti-CD3ε antibody and measured by flow cytometry. The MFI of CD3ε in the GFP+ cells is shown. **(E)** Cells as in (C) were stained with an PE-labeled anti-CD69 antibody and measured by flow cytometry. Cells stained with an isotype antibody serve as control. The MFI of CD69 in the GFP+ cells is shown. (C, D and E) Significance was analyzed by paired t-test (n=3 different donors).

We retrovirally transduced primary T cells, expanded from peripheral blood obtained from healthy human donors, with a construct encoding CD3ε-ERR and GFP. As a control, we used a mock vector encoding for the irrelevant firefly luciferase and GFP (Fig. 6A). As already published ^49^, the surface TCR levels were reduced in the CD3ε-ERR-expressing but not the mock-expressing cells (Fig. 6C and S4A). Importantly and in line with our Jurkat cell-derived data, we found that the total TCR levels were increased in CD3ε-ERR T cells, as quantified by staining CD3ε and TCRαβ (Fig. 6D and S4B). This shows that primary T cells with lower TCR surface levels contain more TCRs in total (i. e. inside) than T cells with physiological TCR surface levels. Hence, our finding based on Jurkat cells was validated in primary human T cells.

### The basal signal correlates with the levels of surface TCRs in primary T cells

Using the TCR retrieval-and-release system, we found that TCRs on the cell surface emit a basal signal that we could measure by quantifying the CD69 expression level (Fig. 5C). To test whether also in primary T cells surface TCRs, but not ER-localized TCRs, would generate a basal signal, we measured CD69 expression in unstimulated CD3ε-ERR- and mock-expressing primary human T cells (Fig. 6E). As expected, the CD3ε-ERR cells with decreased TCR surface levels showed less CD69 upregulation compared to the control. Thus, the magnitude of the basal signal correlated with the amount of TCRs on the surface and not with the total amounts.

As a control, stimulation of TCRs with anti-CD3ε antibodies for 6 hours resulted in a marked upregulation of CD69 compared with the basal, non-stimulated state (Fig. S4C), confirming that CD69 expression serves as a readout of TCR signaling strength also in primary T cells.

Collectively, these findings indicate that TCRs at the cell surface contribute to the basal signaling in primary human T cells, whereas TCRs in the ER do not.

### CD3 phosphorylation and canonical downstream signaling proteins are part of the basal signal

Having established that surface TCRs generate a basal signal, we next sought to characterize the molecular features of this signal. Our TCR retrieval-and-release system provides a unique experimental framework to address this question by enabling a direct comparison between JKεKO KDEL-εRUSH cells lacking surface TCR-mediated basal signaling (without biotin) and cells in which this signaling is restored following biotin-induced TCR trafficking to the plasma membrane. To this end, we identified and quantified phosphorylation sites using mass spectrometry-based phosphoproteomics. We compared the total phosphoproteome in the JKεKO KDEL-εRUSH cells cultured without biotin with the one of cells treated with biotin for one hour. In this unbiased approach we detected a total of 16,884 unique phosphosites (Fig. S5A). In our comparison, 557 phosphosites were upregulated at least 2-fold (log2 = 1.0, green dots, Fig. 7A) and 424 were downregulated at least 2-fold (brown dots). The full dataset is given in supplemental Table S3. Hence, biotin addition and TCR transport to the surface had a clear impact on the phosphorylation events in these cells. Since the expression level of the proteins stayed constant during the hour (Fig. S5B), it was the phosphorylation per protein, i.e. signaling, that changed.

**Figure 7.**
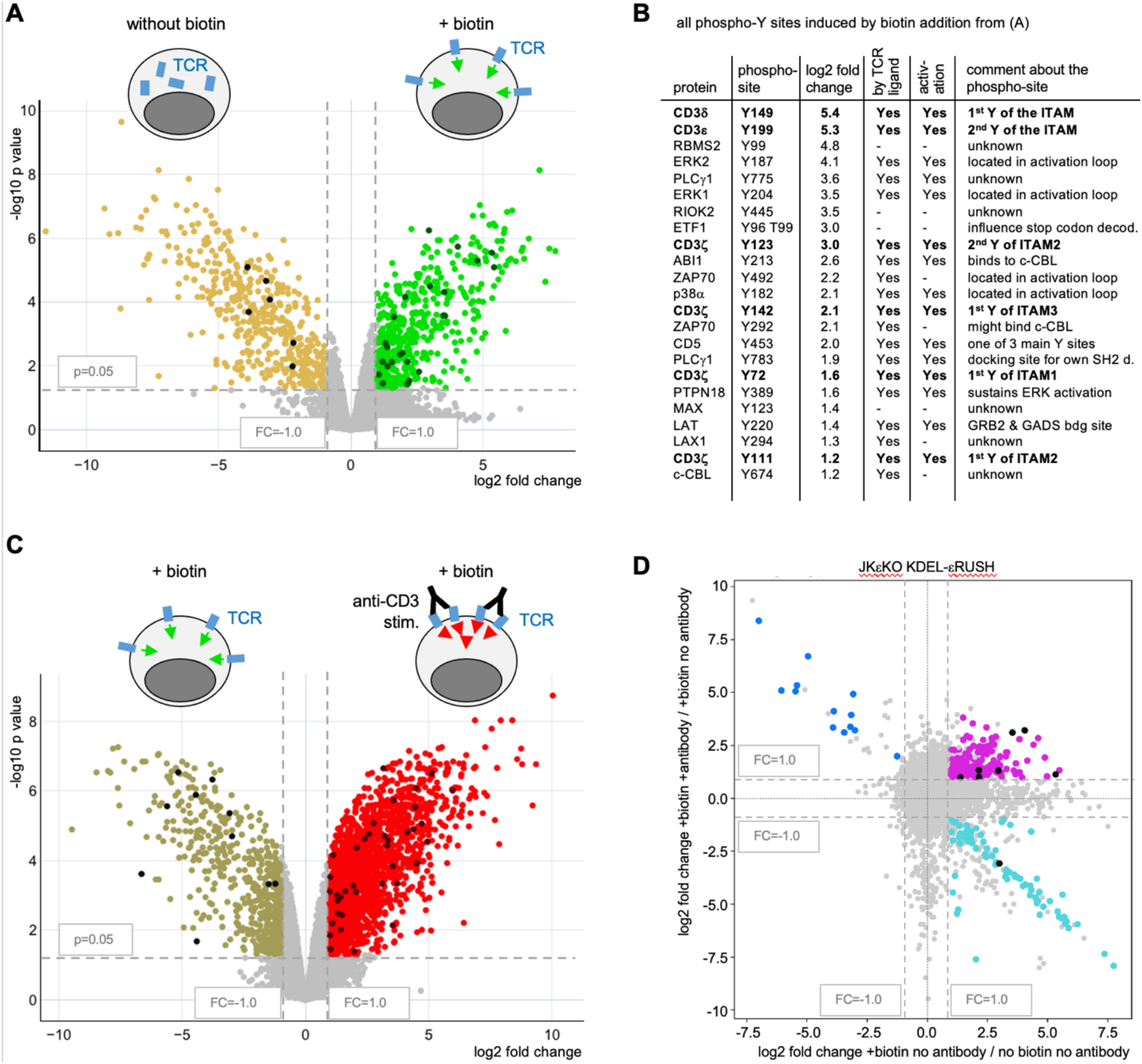
Using the KDEL-εRUSH system to analyze the basal signal with phosphoproteomics. **(A)** JKεKO KDEL-εRUSH cells were cultured for one hour with (+ biotin) or without biotin. After cell lysis, peptides were generated using trypsin. Phospho-peptides were enriched and identified by tandem mass spectrometry. Their intensities in the +biotin condition were compared with those without biotin treatment and visualized as volcano plot based on moderated t-tests. Phosphorylations significantly enriched, p-value <0.05 and log_2_ fold change >1.0, are shown in green and the ones enriched without biotin in brown. The black dots represent Y phosphorylations significantly changed in one condition. **(B)** All phospho-Y peptides enriched in the + biotin sample from (A) are shown. Sites known to be induced by ligand binding to the TCR are indicated in the “by TCR ligand” column and the ones known to have an activatory role in ligand-induced TCR signaling in the “activation” column. **(C)** JKεKO KDEL-εRUSH cells were cultured for one hour with (+ biotin) and left unstimulated or stimulated for 5 min with anti-CD3ε antibodies. Phosphoproteomics and display as in (A) comparing the +/- TCR stimulation conditions. **(D)** Comparison of the basal TCR signal (phosphopeptides from JKεKO KDEL-εRUSH cells with biotin for 1 h versus without biotin, X axis) with the activation signal (phosphopeptides from JKεKO KDEL-εRUSH cells with biotin and stimulated for 5 min with anti-CD3 antibodies versus unstimulated, Y axis). Phosphopeptides significantly enriched both in the basal TCR signature (Table S4) and with the activation signal are shown in purple, those depleted in the basal signal and enriched in the activation signal in blue, those enriched in the basal signal and depleted in the activation signal in cyan, and all others in grey.

Ligand-induced signaling at the TCR is initiated by tyrosine (Y) phosphorylations, such as phosphorylation of the CD3 subunits and early downstream signaling proteins, such as ZAP70, LAT and PLCγ1 ^14^. Focusing on the phospho-Y-containing peptides, we find basal signaling-induced phosphorylation of 23 peptides, including those from CD3ε, CD3δ and CD3ζ (Fig. 7B). This suggests that the basal signal we detected before (Fig. 5 and 6) was initiated by the TCR. Indeed, we also see phosphorylation of downstream proteins, such as ZAP70, LAT, PLCγ1, ERK1 and ERK2 (Fig. 7B). This suggests that the basal signal might use the same signaling machinery as the ligand-induced signal (the latter is called activation signal here, since it leads to the activation of the T cell). We also see serine and threonine phosphorylations of known downstream proteins altered by TCR ligand binding, such as phosphorylation of the adapter NCK1 or the kinase HPK1, as well as dephosphorylation of known downstream proteins, such as the transcription factor NF-AT (supplemental Table S3).

### Identification of the signature of the basal TCR signal

In figure 7A, we compared not only cells having a surface-localized TCR with cells having an ER-localized TCR, but also biotin-treated with untreated cells. Consequently, some of the observed changes could have resulted from the biotin treatment itself rather than from differences in TCR localization. To address this possibility, we compared JKεKO KDEL-εRUSH cells and JKεKO cells, both treated with biotin for one hour. Under these conditions, JKεKO KDEL-εRUSH cells are expected to exhibit the basal TCR signal, whereas JKεKO cells lack a TCR and therefore serve as a negative control. This comparison involves different cell populations cultured on separate plates, and hence numerous differences were detected (Fig. S5C, 8460 up- and 311 downregulated phosphosites). It is not possible to determine which of these reflect basal TCR signaling or simply interculture variability. Nevertheless, all of the phospho-Y changes identified in figure 7A and B were also observed in this comparison (Fig. S5D). These shared changes are therefore unlikely to be a direct consequence of biotin treatment. Concerning the serine and threonine sites, we find 456 phosphorylations that are upregulated in both comparisons, namely with versus without biotin (Fig. 7A) and with versus without TCR (Fig. S5C), see the green sites in figure S6A. We also find 32 phosphorylations being downregulated in both comparisons (Fig. S6A, brown sites). Together with the 23 phospho-Y sites, these changes represent the signature of the basal TCR signal and are listed in Table S4.

Gene Ontology (GO) terms associated with the phosphosites that are upregulated upon induction of the basal signal include “TCR signaling pathway”, “organelle organization”, “cytoskeleton organization” and “intracellular transport” (Fig. S6B and S6C), highlighting that the basal signal is similar to the activation signal and that it might regulate protein trafficking. GO terms of the downregulated phosphosites include “RNA export from the nucleus” and “mRNA transport”, although with low confidence. Anyhow, this could suggest that production of new proteins might be altered by the basal signal.

### The basal signal differs qualitatively from the activation signal

The basal signal could be similar to the activation signal with a quantitative difference, in that the same phosphorylations occur but to lower intensity. In contrast, the basal signal could be qualitatively different from the activation signal, in that some events only occur in one of the two signals.

To obtain insight, we compared the basal TCR signal to the activation signal. We treated the JKεKO KDEL-εRUSH cells for one hour with biotin, to release the TCRs to the cell surface, and then kept the cells unstimulated (basal signal) or stimulated for 5 min with anti-CD3ε antibodies (activation signal). As expected, we see much more changed phosphorylation sites, namely 2097 being up- and 541 being downregulated, as compared to the basal signal (Fig. S6A). Focusing on the TCR, phosphorylation of six Y sites were induced by antibody stimulation (Fig. S7A). Some were already seen in the basal signal and were more phosphorylated upon antibody stimulation (such as CD3ε Y199, CD3ζ Y123 or CD3ζ Y142). Others were only detected in the activation signal (such as CD3ε Y188, CD3ζ Y83 or CD3δ Y160).

Looking at all phosphorylations sites (Y/S/T), we compared the basal signal signature with the activation signal (Fig. 7D). We found that 149 phosphorylations, 7 phospho-Y and 142 phospho-S/T, which are part of the basal signal, were enhanced in the activation signal (Fig. 7D, purple and black sites, and supplemental Table S5). Examples are LAT Y220, LCK S103 and NCK1 S85. 78 other phosphosites were upregulated in the basal signal but disappear upon antibody stimulation (Fig. 7D, cyan sites), such as NCK1 S91 and Kidins220 S1681. These events are specific for the basal signal. This differential phosphorylation of NCK1 is interesting, since NCK1 binds to the TCR and has been implicated in ligand-induced TCR signaling ^51,52^. Finally, 13 phosphosites were downregulated by the basal signal and then induced upon antibody stimulation (Fig. 7D, blue sites), such as TCF20 S1361.

Together this indicates, that while some signaling events are similar in basal versus activation conditions, others are specific for the basal or for the activation signal. Some of these differences might already originate at the TCR, since they include CD3 subunit phosphorylations.

### LCK-dependent phosphorylation is part of the basal TCR signal

It was suggested that the basal signal in T cells is independent of the presence of a TCR ^30^. This is in contrast to our findings on the nature of the basal signal that includes Y phosphorylations of the CD3 subunits (Fig. 7B and S5D). These phosphorylations are mediated by the Src family kinase LCK ^15^. Hence, our prediction was that LCK’s kinase activity was required for the generation of the basal signal. To test this, we quantified CD69 expression using flow cytometry and JKεKO KDEL-εRUSH cells treated or not with biotin as in figure 5C. The control cells incubated with DMSO upregulated CD69 after 7 and 22 hours of biotin treatment (Fig. 8A), confirming our previous results on the induction of the basal signal (Fig. 5C). Incubation of the cells with the Src kinase inhibitor PP2, which inhibits LCK ^53^ abrogated this basal signal (Fig. 8A), indicating that LCK activity was required. As a control, biotin treatment of JKεKO cells, which do not possess a TCR, did also not lead to CD69 upregulation (Fig. 8B). This confirms that a TCR is required for the generation of the basal signal and that biotin itself does not lead to CD69 upregulation.

**Figure 8.**
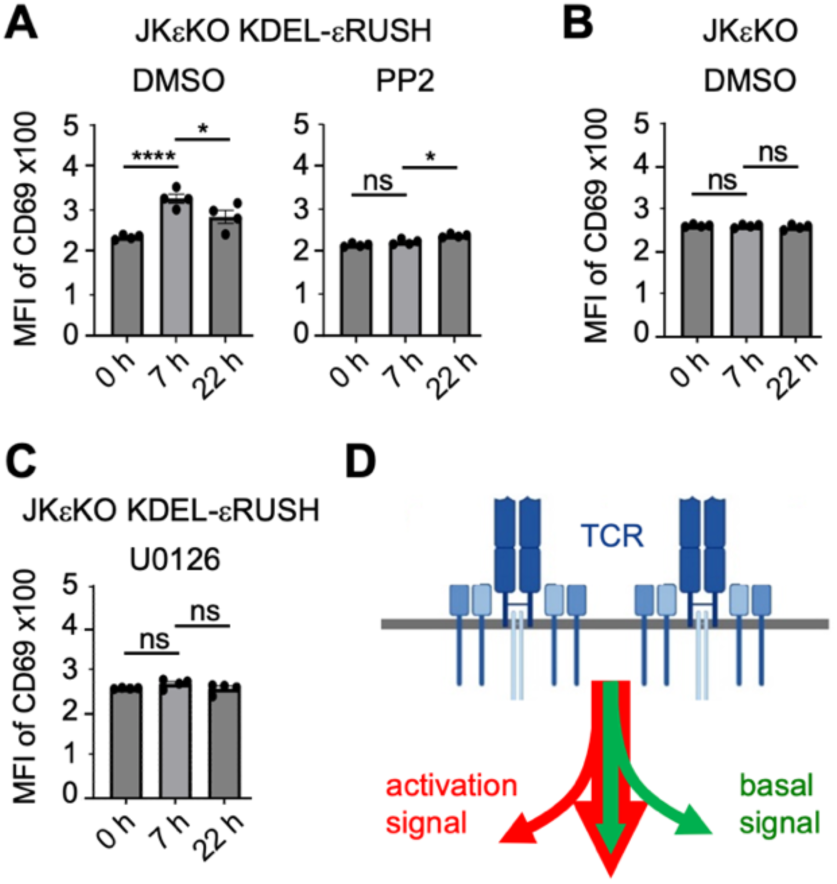
LCK and ERK1/2 are part of the basal signal. **(A)** JKεKO KDEL-εRUSH cells were cultured in medium without biotin and incubated with the vehicle DMSO or with 10 µM of the Src kinase inhibitor PP2. In addition, cells were treated for 7 or 22 hours with 40 µM biotin or were kept untreated (0 h). Then cells were stained with an APC-labeled anti-CD69 antibody and measured by flow cytometry. The MFI of CD69 is shown. Significance was analyzed by two-way ANOVA (n=3; shown one representative experiment in quadruplicates). **(B)** The experiment using DMSO was performed and analysed as in (A) with JKεKO cells. **(C)** The experiment was performed and analyzed as in (A) incubating the cells with 20 µM of the MAP kinase inhibitor U0126. **(D)** The basal signal (green) is similar to the activation signal (red), but with reduced intensity (vertical arrows). In addition, basal-specific and activation-specific signaling events exist (curved arrows).

In T cells the RAS pathway, including the MAP kinases ERK1 and ERK2, is downstream of ligand-induced TCR signaling and contributes to CD69 expression ^54^. We detected phosphorylation of ERK1 and ERK2 as part of the basal signal signature (Fig. S5D and Supplemental Table S4). To assess their contribution to this signal, we inhibited these MAP kinases with the inhibitor U0126. Indeed, upon U0126 incubation, biotin treatment of JKεKO KDEL-εRUSH cells no longer induced CD69 upregulation, demonstrating that ERK1 and ERK2 are part of the basal signal. The finding that LCK and ERK1/2 are part of the ligand-independent basal signal, is in line with the idea that the basal signal is similar to the activation signal, although at lower intensity.

## Discussion

In this study we established two systems with which TCRs and assembly intermediates are kept inside the T cell and released to the cell surface upon biotin addition. In one system these complexes are retained in the ER (Ii-εRUSH) and in the other one they are constantly retrieved back into the ER from post-ER compartments (KDEL-εRUSH). Our data suggest that (i) TCR assembly is only completed in post-ER compartments, and (ii) TCRs only emit a ligand-independent basal signal from the cell surface and not from the ER. This has enabled us to get hold on the basal signal and identify its signature.

Both, the Ii-εRUSH and the KDEL-εRUSH systems, rely on a fusion protein composed of CD3ε with its ectodomain being attached to SBP and GFP, called εRUSH. We have chosen this arrangement, because large domains fused to the ectodomain of CD3ε do not interfere with the assembly or the function of the TCR ^34^. As expected, εRUSH assembles with the other TCR’s subunits. Further, this arrangement leaves the intracellular signaling domains unobstructed. Indeed, εRUSH-containing TCRs at the cell surface were signaling competent after antibody stimulation.

We expected that full TCR assembly would occur in the ER, because the full αβεδεγζζ TCR was formed when all subunits were *in vitro* translated into B cell-derived ER vesicles ^12^. However, by retaining εRUSH in the ER using the Ii-εRUSH system, only partial complexes formed. Consequently, upon release by biotin addition they failed to reach the cell surface. Likewise, in TCR subunit-deficient T cell lines, the partial complexes did not reach the cell surface ^11,55–58^. After longer times of biotin treatment more TCRs reached the cell surface in the the Ii-εRUSH system. We propose that these were newly synthesized TCRs formed in the presence of biotin and thus were never retained in the ER, allowing their assembly to occur in the ER as well as in post-ER compartments. When we performed an intracellular flow cytometry stain that detected folded CD3ε molecules, i.e. folded εRUSH, there was a clear increase after 15 min of biotin-mediated release in the Ii-εRUSH system. This indicated that some εRUSH molecules only folded correctly upon release from the Ii-hook and (transient) transport to post-ER compartments. This enhanced folding might have been assisted by pairing with CD3δ or CD3γ ^39,40^. Indeed, we observed increased assembly of TCRα, TCRβ and CD3δ upon release of εRUSH from the Ii-hook.

We also used the KDEL-hook with which the retained εRUSH is kept inside the cell by leaving the ER and constantly being retrieved back into the ER ^37^. In this case TCR assembly could occur in the ER and in post-ER compartments and indeed the complete αβεδεγζζ TCR complex formed during this retrieval. Consequently, after release, TCRs reached the cell surface within 30 min and these were the pre-formed ones that assembled prior to biotin addition. There was no increase in εRUSH folding after release, suggesting that proper folding and pairing with CD3δ or CD3γ had taken place during retrieval.

In conclusion, some steps of TCR assembly cannot occur in the ER, but in compartments that are visited by the KDEL retrieval sequence, such as the ER-Golgi intermediate compartment (ERGIC) or the cis-Golgi ^59^. We suggest that different chaperones, a different glycosylation of TCR subunits or a different lipid environment might enable complete TCR assembly in post-ER compared to ER compartments. Further, the fact that the Ii-hook is a membrane-bound protein compared to the soluble KDEL-hook might have influenced εRUSH folding and assembly.

Upon synchronous release of TCRs from the ER in the KDEL-εRUSH system, TCRs transiently accumulated at the surface, followed by a reduction in surface TCR levels over time. The total cellular TCR abundance, comprising both intracellular and surface-localized receptors, was highest when TCRs were retained intracellularly and decreased upon biotin addition. To obtain mechanistic insight into these dynamics, we developed a mathematical model capturing the main steps of TCR production, transport and degradation. Our quantitative data from the KDEL-εRUSH system, amounts of surface and total TCRs as well as the proportions of post-ER and ER-localised TCRs, required two feedbacks by surface TCRs: a slowed export from the ER to the surface and enhanced recycling. In contrast, feedback from ER-localized TCRs was not required to reproduce the experimental observations. These findings suggest that surface TCRs generate a signal independently of ligand binding, which we refer to as the ligand-independent basal signal. Ligand-induced receptor signaling has been extensively studied, facilitated by the ability to compare cellular signaling states before and after ligand addition. In contrast, ligand-independent basal receptor signaling remains poorly understood. A major challenge is that basal signaling occurs constitutively, making it difficult to establish an appropriate reference state in which basal receptor signals are absent. Genetic knock-outs are limited by compensatory cellular adaptations to the loss of the protein ^60,61^. Pharmacological inhibition of receptors is often not possible due to the lack of a suitable inhibitor, and inhibition of downstream signaling proteins may interfere with signaling mediated by multiple receptors, thereby confounding the interpretation of the results.

Here, we introduce a new approach to study basal signaling by the TCR. Our strategy involves retaining TCRs in the ER, where basal signaling is absent, and subsequently releasing them to the cell surface, where basal signaling occurs. Thus, we can compare two different states to get hold on the basal signal. We provide experimental evidences that basal signaling originates from surface-localized, but not ER-localized, TCRs.

i. phosphoproteomic analysis identified 981 differential phosphorylation events between biotin-treated (surface TCRs) and untreated (ER TCRs) JKεKO KDEL-εRUSH cells. Of these, 511 were caused by TCR trafficking rather than the biotin treatment itself. These phosphosites constitute the basal signal signature (supplementary Table S4).
ii. expression of CD69, a downstream marker of ligand-induced TCR signaling ^62^, was slightly but significantly increased upon TCR release from the ER. Since biotin alone did not cause this increased expression, it was the localization of the TCR.
iii. the TCRα mRNA, called TRAC, is a hallmark of a proposed TCR-independent basal signal that engages the downstream TCR signaling machinery, including LAT ^30,31^. TRAC expression was higher in cells with surface-localized TCRs than in cells with ER-retained TCRs.

Together this shows that surface TCRs produce a basal signal that ER-localized TCRs do not. Firstly, we see phosphorylation of CD3 as being part of the signal. Secondly, the signal as measured by CD69 upregulation disappeared when CD3 phosphorylation was inhibited by the Src kinase inhibitor PP2. Hence, TCRs are the origin of the basal signal. This contrasts with an earlier study suggesting that basal signaling is independent of the TCR ^30^. The authors detected basal signaling in Jurkat T cells lacking TCRα or TCRβ expression. Although these cells lack surface TCRs, they express CD3ζat the surface ^58^, raising the possibility that surface CD3ζ might have mediated the observed basal signal.

How can TCRs signal without ligand binding? According to the allostery model, TCR signaling starts when the TCR adopts the active conformation ^18,63^. Even in resting T cells, some TCRs spontaneously adopt this conformation and may continuously cycle between inactive and active states, with the latter being signaling-competent even without ligand binding ^22,64^. (Swamy *et al*, 2016). Additionally, TCRs may form ligand-independent clusters at the cell surface ^3,65^. Hence, both criteria for TCR signaling - active conformation and clustering ^2^ - may be met without ligand binding. Consistent with this, spontaneous TCRs phosphorylation can be detected using pervanadate, which inhibits phosphatases ligand-independently (Secrist et al, 1993), or by increasing the proportion of TCRs that are spontaneously in the active conformation ^22^.

ER-localized TCRs might not support basal signaling because LCK, the kinase LCK that phosphorylates CD3, is anchored to the plasma membrane by post-translational palmitoylation and myristoylation ^67–70, 68,71^(Bijlmakers *et al*, 1997; McCabe & Berthiaume, 1999; Resh, 1999)limiting its access to ER-localized TCRs for phosphorylation. Furthermore, spontaneous TCR clusters are driven by cholesterol and sphingomyelin ^72,73^, which are scarce in the ER but abundant at the cell surface ^74^. Hence, signaling-promoting TCR clusters may form at the plasma membrane but not in the ER.

Our signature of phosphorylation events that occur in basal TCR signaling show that the basal signal (Fig. 8D, green arrows) is similar to the ligand-induced activation signal (Fig. 8D red), but with reduced intensity (vertical arrows). This relates to many classical downstream signaling proteins of the TCR and includes some CD3 phosphorylation events. Indeed, it was suggested earlier that the basal signal might be very similar to the ligand-binding induced signal, but at lower in intensity ^30^. In addition, basal-specific and activation-specific signaling events also exist (Fig. 8D, curved arrows, and supplemental Table S5).

A difference between ligand-independent basal and ligand-induced activation signals is the different phosphorylation of the CD3 ITAMs. (Pettmann *et al*, 2021; Yousefi *et al*, 2019)Some sites are phosphorylated in the basal TCR signal and not further enhanced by ligand-binding to the TCR, Other sites are phosphorylated in the basal TCR signal and stay constant upon ligand-binding (Fig. S7). This might be explained by different thresholds controlling the phosphorylation of these sites. Interestingly, different patterns of ITAM phosphorylation might include mono-phosphorylation. In contrast to double phosphorylation which recruits the kinase ZAP70 and delivers an activation signal, mono-phosphorylated CD3 might recruit inhibitory signaling proteins ^75–77^. The latter most likely dampens the generation of a full activation signal and would explain why basal signaling does not trigger T cell activation.

Overall, our study establishes a novel conceptual and experimental framework for investigating receptor assembly, trafficking and basal receptor signaling.

## Methods

### Plasmids

The Ii-Str_SBP-EGFP-Ecadherin (Addgene plasmid #65288) and Str-KDEL_TNF-SBP-EGFP (Addgene plasmid #65278) plasmids ^32^ were purchased from Addgene. All plasmids generated in this study were generated using standard molecular cloning techniques, like polymerase chain reaction (PCR), restriction enzyme digestion and ligation or Gibson assembly ^78^. The plasmids are listed in Table S1 together with the corresponding coded protein and the cloning strategy. All used oligonucleotides are listed in Table S2. All generated plasmids were verified by colony PCR and Sanger sequencing.

### Cell lines

The Jurkat (JK) T cell line is derived from a human T cell leukemia ^79^. JK cells knocked out for CD3ε (JKεKO) expression were generated by CRISPR/Cas9 knock out for the respective TCR subunit ^34^. JK, JK knock out (KO) cell lines and all generated JK cell lines were cultured in biotin-containing RPMI 1640 medium (Anprotec) supplemented with 10% FBS, 10 mM HEPES, 100 U/ml penicillin and 100 µg/ml streptomycin and 0.1% β-mercaptoethanol (β-MeOH, Sigma), further referred to RPMI medium, at 37°C and 5% CO_2_. Alternatively, cells were washed in PBS and cultured in biotin-free RPMI 1640 medium (USBiological) supplemented with 10% biotin-free FBS, 10 mM HEPES, 100 U/ml penicillin and 100 µg/ml streptomycin and 0.1% β-MeOH, further referred to medium without biotin, at 37°C and 5% CO_2_. Biotin-free FBS was generated by overnight incubation of FBS with streptavidin-sepharose beads and subsequent sterile filtering through a 0.22 µM filter. HEK293 T cells (CRL-1573, from ATCC) were cultured in DMEM medium (Pan Biotech) supplemented with 10% FBS, 1 mM sodium pyruvate (Anprotec), 100 U/ml penicillin and 100 µg/ml streptomycin at 37°C and 7.5% CO_2_.

### Lentiviral production and transduction

Lentiviral production was done as described previously ^80^. The concentrated lentiviral particles were resuspended in PBS and frozen in aliquots at -80°C. Lentiviral particles were produced from the plasmids pCJ-02, and -035, see supplement Table S1 and S2.

For lentiviral transduction JK and JK KO cells were adjusted to 1x10^6^ cells/ml in RPMI medium and plated in a 24-well plate. Cells were transduced by addition of the lentivirus and soft resuspension. After 24 hours new RPMI medium was added and after 72 hours cells were washed in PBS and resuspended in medium without biotin or kept in RPMI medium. Transduction of cells was monitored by GFP fluorescence and TCR surface expression was analyzed by flow cytometry. JKεKO cells transduced with pCJ-02 vector are called JKεKO Ii-εRUSH cells and JKεKO cells transduced with pCJ-035 vector are called JKεKO KDEL-εRUSH cells. Both cell lines were sorted according to GFP fluorescence and TCR surface expression in RPMI medium or in medium without biotin after addition of biotin for 60 min. After sorting, cells were cultured in medium without biotin as well as in RPMI medium and used for all experiments.

### Flow cytometry

Cells were stained to detect surface proteins according to standard protocols. Briefly, cells were washed with FACS buffer (PBS with 2% FBS), and incubated for 20 min on ice with the desired antibody dilutions in FACS buffer. Afterwards, the cells were washed and resuspended in FACS buffer. If cells were stained with an unlabeled antibody, cells were washed and stained again with a fluorescently labeled secondary antibody, washed, and resuspended in FACS buffer. Cells were measured by an Attune NxT flow cytometer and data were analyzed with FlowJo software version 10.0.7r2. The antibodies used in this study were: AF488-labeled anti-CD3ε (clone UCHT1; BioLegend), BV421-labeled anti-CD3ε (clone UCHT1; BioLegend), AF647-labeled anti-CD3ε (clone UCHT1; BioLegend), PE-labeled anti-CD3ε (clone UCHT1, eBioscience), PE-labeled anti-Vβ8 (BD Biosciences), goat anti-GFP (Rockland), AF680-labeled anti-goat IgG (Invitrogen), APC-labeled anti-CD69 (Invitrogen), AF647-labeled anti-GFP (Invitrogen).

Cells were stained to detect intracellular proteins according to standard protocols. Briefly, cells were fixed with 2% PFA in FACS buffer for 30 min on ice. Subsequently, cells were washed and permeabilized with 87.7% methanol for 30 min on ice. Then, cells were washed and stained for 2 h at room temperature with the desired antibodies. Finally, cells were washed and resuspended in FACS buffer. Cells were measured and analyzed as above. The antibodies used in this study were: BV421-labeled anti-CD3ε (clone UCHT1; BioLegend), and AF647-labeled anti-HA.11 (BioLegend).

### Time course of biotin addition

Cells were cultured in medium without biotin and left untreated or treated for different times with 40 µM D-biotin (Sigma) or were cultured in RPMI medium at 37°C, 5% CO_2._ If cells were treated with puromycin, 1 μg/ml puromycin (Sigma) was added 30 min before addition of biotin and puromycin was kept in culture. Afterwards, cells were washed, stained, and analyzed by flow cytometry.

### CD69 upregulation

Cells were cultured in medium without biotin and left untreated or treated for 1 or 16 hours with 40 µM biotin or cultured in RPMI medium at 37°C, 5% CO_2_. When mentioned, cells were pre-incubated either in the presence of 10 µM PP2, 20 µM U0126 or DMSO 10 min prior to the incubation with biotin. Then cells were left unstimulated or stimulated with 5 μg/ml anti-CD3ε (clone UCHT1) for 6 h at 37°C, 5% CO_2_ and biotin was kept in culture. Afterwards, cells were washed, stained and analyzed by flow cytometry as described above.

### Live-cell imaging

Eight-well glass chambers (Lab-Tek II chambered coverglass #1.5 borosilicate, Nalgen Nunc International) were cleaned with hot water, DI water, ethanol and DI water (three times each). Air-dried chambers were plasma cleaned (HPT-100, Harric Plasma) and coated with 0.1 mg/ml biotinylated poly-L-lysine (bioPLL). bioPLL was coupled at 50:1 ration of NHS-biotin to 70 kDa poly-L-lysine (10 mg/ml) in PBS pH7.8 and dialyzed into water. Surfaces were blocked with 0.5% BSA and sequentially coated with 25 µg/ml streptavidin and 5 µg/ml biotinylated ICAM1 in HEPES-buffered saline solution (HBS, 20 mM HEPES pH7.4 + 150 mM NaCl). Glass chambers were washed and stored in HBS, which was exchanged with 200 µl of 2x adherence solution (Hanks buffered saline solution (HBSS) with 2 mM EGTA, 10 mM MgCl_2_) before cells were added.

Prior to imaging 0.5 × 10^5^ cells were pelleted at 250 g, resuspended in 200 µl HBSS and added to coated glass chambers containing 2x Adherence solution equilibrated to 37°C. Cells were incubated for 30 minutes for adhering before adding biotin (CarboSynth) to a final concentration of 40 µM. Images were acquired on a custom-build inverted microscope platform (Visitron Systems GmbH, Nikon Instruments and Cairn Europe) with a 60x objective (CFI APO TIRF oil immersion NA1.49, Nikon) using widefield microscopy. Image acquisition and deconvolution was performed using VisiView software (Visitron Systems GmbH), and images were further processed using ImageJ software.

### Immunoprecipitation

Cells were cultured in medium without biotin and left untreated or treated with 40 µM biotin for different times at 37°C, 5% CO_2_. Then cells were lysed in lysis buffer containing 20 mM Tris-HCl pH 8, 137 mM NaCl, 2 mM EDTA, 10% glycerol, protease inhibitor cocktail, and 0.5% Brij96V for 30 min at 4°C. Subsequently, nuclei and insoluble material were removed by centrifugation. The cleared lysate was incubated with 10 μl protein A-coupled Sepharose and 1 μg of the goat anti-GFP (Rockland) antibody for 4 h at 4°C. After washing, the immunoprecipitated proteins were separated by non-reducing or reducing sodium dodecyl sulfate polyacrylamide gel electrophoresis (SDS-PAGE) and Western blotting was performed.

### SDS-PAGE and Western blotting

Purified proteins and cell lysates were separated by 10% or 12% non-reducing or reducing SDS-PAGE. Proteins were transferred to PVDF membranes by wet transfer. After blocking with 5% milk in PBS containing 0.1% Tween 20, the membranes were incubated with the primary antibodies followed by incubation with horseradish peroxidase-conjugated secondary antibodies. Proteins were detected using an Image Quant LAS 4000 Mini (from GE Healthcare Life Sciences). The antibodies used in this study were: anti-CD3γ (Epitomics), anti-CD3δ (Santa Cruz), anti-CD3ζ, anti-TCRα (Santa Cruz), anti-TCRβ (Santa Cruz), anti-GFP (Roche), anti-HA (Millipore), anti-streptavidin (Rockland), anti-pERK (Cell Signaling) and anti-GST (Bethyl).

### RT-qPCR

JKεKO KDEL-εRUSH cells were cultured in biotin-free medium and either left untreated or treated with 40 µM biotin for 1, 6, or 24 h. Cell pellets were lysed for RNA extraction using the RNeasy Extraction Kit (Qiagen). cDNA was synthesized from 1 µg of freshly isolated RNA, diluted 1:10 in nuclease-free water, using the First Strand cDNA Synthesis Kit (Fisher Scientific). RT-qPCR was performed using the FastStart Universal SYBR Green Master Kit (Sigma). TRAC expression was analyzed using gene-specific primers (Sigma), with β2 microglobulin (B2M) serving as a housekeeping gene for normalization (Table S2). Each sample was analyzed in technical triplicates. Amplification was performed using a Bio-Rad 384-well real-time PCR system, and quantification was based on the resulting quantification cycle (Cq) values.

### Generation and analysis of the CD3ε-ERR-expressing primary human T cells

Primary human T cells were generated from peripheral blood obtained from healthy donors after informed consent, under protocols approved by the Ethics Committee of the Royan Institute, and were activated, retrovirally transduced, and expanded as previously described ^49^. Cells were transduced with either SFG-CD3ε-ERR-IRES-eGFP or the control SFG-FFluc-IRES-eGFP vector. The effects of CD3ε-ERR expression on the TCR were assessed by surface and intracellular staining with anti-CD3ε -APC (clone OKT3; Biolegend) and anti-TCRα/β-PerCP/Cyanine5.5 (clone IP26; Biolegend), followed by flow-cytometry analysis. For functional assessment, CD3ε-ERR and control mock T cells were stimulated with plate-bound anti-CD3 (clone OKT3; Miltenyi Biotec), as previously described ^49^. After 18 h, T cell activation was quantified by surface expression of CD69 using anti-CD69-PE (clone FN50; BioLegend).

### Sample preparation for phosphoproteomics

6x10^6^ cells were incubated overnight in medium without biotin at 37°C and 5% CO_2_. Then biotin (or H_2_O as control) was added to the cells to a final concentration of 40 µM and incubated for 60 min at 37°C and 5% CO_2_. Then, cells were either left unstimulated or stimulated with 10 µg/ml anti-CD3 antibodies (clone UCHT-1 and OKT3, each 5 µg/ml) for 5 min. Subsequently, cells were washed with ice-cold PBS and lysed in 100 µl lysis buffer containing PBS, 10 mM HEPES, 2% SDS (pH 7.4) and 1x HALT^TM^ Protease and Phosphatase Inhibitor Cocktail (Thermo Fisher Scientific) at 37°C for 10 min. Lysates were sonicated with a homogenizer (Bandelin Sonopuls mini20) at 65% intensity with 2 cycles of 4 pulses per 10 s. The sonicated lysates were centrifuged for 20 min at 20.000 g. Protein concentration in the cleared lysate was determined with the Pierce^TM^ BCA Protein Assay Kit (Thermo Fisher Scientific). Per sample, 100 µg of protein was used for further phosphoproteomic analysis. The replicates per condition were: JKεKO + biotin, 5 replicates; JKεKO KDEL-RUSH w/o biotin, 5 replicates; JKεKO KDEL-RUSH + biotin, 4 replicates; JKεKO KDEL-RUSH + biotin + anti-CD3ε-antibodies, 4 replicates.

Protein samples were treated with 0.3 µl Benzonase® Nuclease (Merck-Millipore) and incubated for 30 min at room temperature to remove residual DNA. Protein purification, digestion and phosphopeptide enrichment protocols were performed in 96-well plates (1 mL, Agilent Technologies) using a STARlet liquid handling system (Hamilton, Bonaduz, Switzerland) equipped with a 96-well magnet plate (Magnum Flex, Alpaqua). Proteins were incubated with shaking with 10 mM DTT for 30 min at 65°C followed by 50 mM chloroacetamide (CAA) for 30 min at 22°C. 10 μl of 50 mg/ml paramagnetic SP3 beads (Cytiva Europe) were added followed by incubation in 80% ethanol (EtOH) for 20 min at room temperature. Samples were washed three times with 80% EtOH and resuspended in 50 μl 100 mM ammonium bicarbonate, pH 8.5, containing trypsin/lys-C mix (Promega V5073) in a 1:100 (w/w) protease to protein ratio for overnight digestion. After 16 h at 37°C the supernatant without beads was transferred into new wells and digestion was stopped by adding 30 μl of 5% formic acid (FA). Peptides were desalted using 96 well PurePepBroad plates (Affinisep) pre-activated with MeOH, pre-washed with 80% MeCN, 0.1% FA and pre-equilibrated two times with 0.1% FA. Peptides were loaded into 100 µl 0.1% FA and washed two times with 400 μl 0.1% FA. Peptides were collected using two subsequent elutions with 100 μl 80% MeCN, 0.1% TFA into a collection plate. 5% of each sample was collected into a low binding 96 well plate for protein-level analysis as input and dried by vacuum centrifugation. For phosphopeptide enrichment, peptides in 200 μl binding buffer (80% MeCN, 0.1% TFA) were incubated with 5 µl Fe-IMAC beads (Cube Biotech, 31501-Fe) for 20 min at room temperature according to manufacturer’s instructions, washed three times with 200 μl of 80% MeCN, 0.1% TFA and eluted from the beads using 2 x 50 μl freshly prepared 50% MeCN, 2.5% ammonia. The supernatant was transferred to a low binding 96 well plate, acidified by addition of 30 μl 5% FA, and dried by vacuum centrifugation. Phosphopeptides and input samples were loaded onto Evotips (EvoSep), according to manufacturer’s instructions.

### Mass spectrometry analysis

For liquid chromatography mass spectrometry (LC-MS) analysis, an Evosep One system (Evosep) was coupled online to an Exploris 480 mass spectrometer (Thermo Fisher Scientific). Peptides were separated using a 68-minute gradient (Whisper Zoom 20 SPD workflow, Evosep) with a nano-LC column (Aurora Elite column 15 cm x 75 µm, AUR3-15075C18, Ionoptics) in a column oven at 50°C. Data were acquired in data-independent acquisition (DIA) mode. Each acquisition cycle consisted of one full MS survey scan (40% RF lens, 300% normalized AGC target, 45 ms maximum injection time, 350–1400 *m/z*, 120k resolution) followed by MS/MS fragment spectra (50% RF lens, 1000% normalized AGC target, 54 ms maximum injection time, 30k resolution) acquired by higher-energy collision-induced dissociation (HCD) at a normalized collision energy of 28%. Fragment ions were collected across three precursor *m/z* ranges: from 361 – 450 with 17 *m/z* isolation width, from 450 – 800 with 9 *m/z* isolation width and from 800 – 1100 with 17 *m/z* isolation width. Isolation windows overlapped by 1 m/z.

For protein and peptide identification and quantification, a library free search was performed using FragPipe version 23.1 ^81^ implemented with DIA-NN version 1.8.2 beta 8 ^82^ against the UniProt proteome set for *Homo sapiens* (release 2023-05). Trypsin was set as protease with 2 allowed missed cleavages. Carbamidomethylation of cysteine was selected as fixed modification, N-terminal excision of methionine and oxidation of methionine were selected as variable modifications with a maximum number of 3. For the phosphopeptide enriched samples additionally phosphorylation at serine, threonine and tyrosine was set as variable modification and the maximum number was increased to 5. For precursor ion generation, peptides ranging from 7 to 45 amino acids, precursor charge from 2 to 4 and precursor mass range from 300 to 5000 were used. The protein group and precursor output files from DIA-NN were used. For quantitative proteome analysis, all proteins with non-zero label-free quantification (LFQ) intensities in 4 out of 4 or 5 replicates in both control and treated group were selected for further analysis. Missing values were imputed by sequential imputation using the R function impSeqRob. For phosphoproteome analysis, identifications of phosphorylated peptides with non-zero label-free quantification intensities in 3 out of 4 or 5 replicates in either control or treated group and with a localisation probability of >=0.75 were retained, log transformed, quantile normalized and data for phosphopeptides with 3 or more missing values in one group were imputed by left-censored missing value imputation using the R function impute.MinProb. For analysis of differential abundance, log_10_ transformed intensities were subjected to a moderated t-test using linear modelling implemented in the R package limma ^83^ with correction of p-values for multiple testing.

### Gene Ontology (GO) terms

GO enrichment analyses were performed using G:Profiler ^84^, version e114_eg62_p19_27110d83, database updated on 20/03/2026, through its API and Python client library.

### Statistics

Data are represented as mean ± SD. All differences between experimental groups were analyzed with the unpaired or paired t test, 1-way ANOVA or 2-way ANOVA using GraphPad Prism (v8.4.3; GraphPad Software). Significant differences were considered when the p values were less than 0.05. (ns = non-significant, *p < 0.05, **p < 0.01, ***p < 0.001, and ****p < 0.0001). n refers to the number of independently performed experiments.

## Supporting information

Supplement: Mathematical Modeling

## Acknowledgments

We thank Sascha Yousefi for providing plasmids and helping with cloning, Angela Naumann from the Life Imaging Center (LIC) for her support in live-cell imaging, Kerstin Fehrenbach for technical assistance and Nicole Gensch, Pavel Salavei and Johannes Kaiser from the BIOSS Signaling Factory for their support. This work was supported by the German Research Foundation (DFG) through the Excellence Initiative of the German Federal and State Governments BIOSS EXC-294 and CIBSS EXC-2189 (project 390939984) and the SFB1381 (Project ID: 403222702). CJ and SH were supported by the DFG through GSC-4 (Spemann Graduate School). VT was funded by the Hans A. Krebs Medical Scientist Programme, Faculty of Medicine, University of Freiburg and the Baden-Württemberg Stiftung (‘ArrhythMEC’).

## Author contributions

Conceptualization, WWS; T cell and ER retention/retrieval experiments, CJ, SH and GF; development and validation of mathematical model, SB and VT; primary T cell experiment, HE and MB; live-cell imaging, CJ, DP and BL; cloning strategy, CJ and AM; RT-qPCR, EM and FS, phosphoproteomics and analysis, SH, KR, MD, FD and PH; writing of the manuscript, CJ, VT and WWS; intellectual input, CJ, SB, FD, PH, BA, VT, and WWS; funding acquisition, WWS, VT, MR and JT.

## Supplementary Material

Supplementary Material is available: 7 supplemental figures and 2 tables as one PDF document, 2 movies as avi files, the mathematical model as one PDF document, and 3 tables as Excel files.

