## Supplementary Figures and Tables S1 and S2 for "An ER Retrieval-and-Release System Reveals Ligand-Independent Basal Signaling by Cell Surface TCRs"

Claudia Juraske *et al.*

**Content:** page

| | | |
|---|---|---|
| Figure S1 | Analysis of the Ii-εRUSH system in JKεKO and JK WT cells | 2 |
| Figure S2 | Analysis of the KDEL-εRUSH system in JKεKO and JK WT cells | 3 |
| Figure S3 | Kinetics of TCR surface expression in the Ii-εRUSH and KDEL-εRUSH systems | 4 |
| Figure S4 | Analysis of primary human T cells expressing CD3ε-ERR | 5 |
| Figure S5 | Analysis of the phospho-proteomics data of the basal TCR signal | 6 |
| Figure S6 | Identification of the basal TCR signal | 7 |
| Figure S7 | Analysis of the phospho-proteomics data of the activation TCR signal | 8 |
| Table S1 | List of plasmids | 9 |
| Table S2 | List of oligonucleotides | 10 |

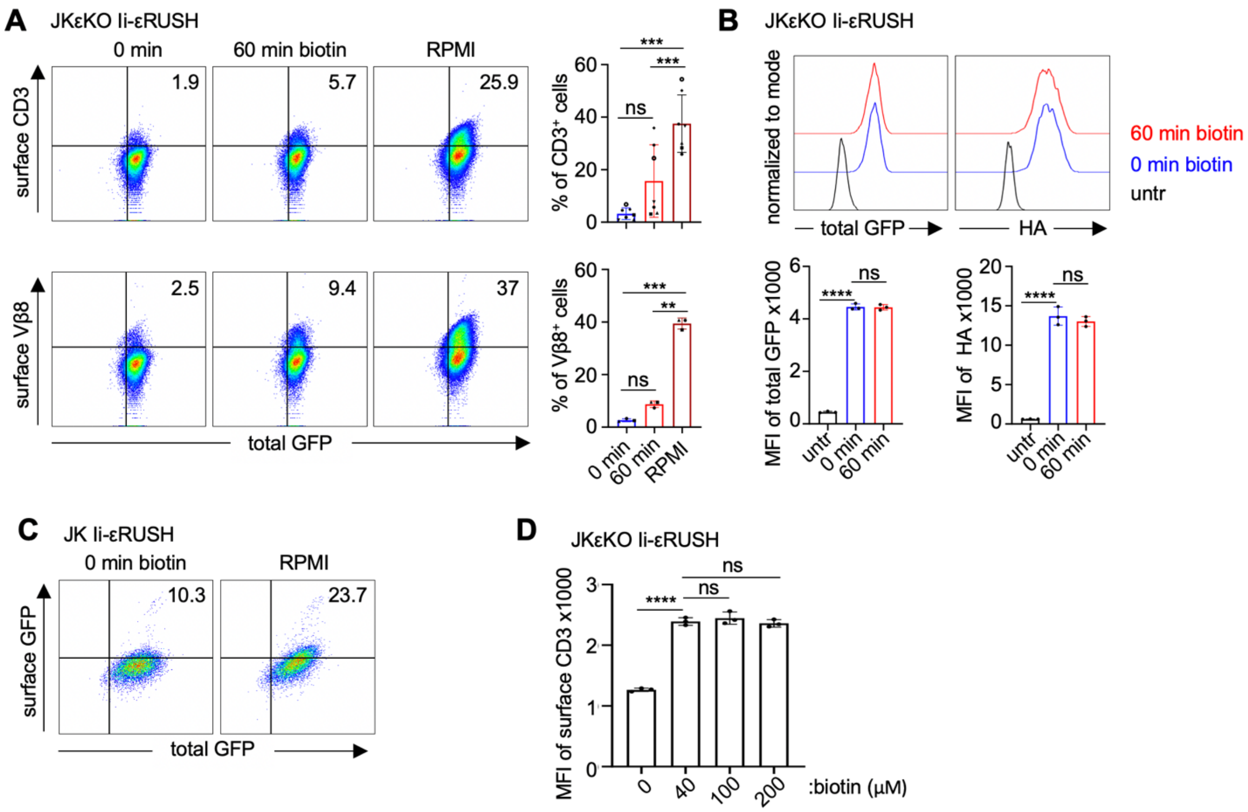


**Figure S1. Analysis of the Ii-εRUSH system in JKεKO and JK WT cells**

**(A)** JKεKO Ii-εRUSH cells were cultured without biotin and left untreated (0 min) or treated for 60 min with 40 μM biotin or cultured in biotin-containing RPMI medium (RPMI). Cells were stained with an AF647-labeled anti-CD3ε (top panels) and PE-labeled anti-Vβ8 antibodies (lower panels), and TCR surface levels were measured by flow cytometry. Percentages of surface $CD3^+$ and surface $TCR\beta^+$ within the $GFP^+$ cells are plotted. Significance was analyzed by ordinary one-way ANOVA (n=3).

**(B)** JKεKO Ii-εRUSH cells were cultured without biotin and left untreated (0 min) or treated for 60 min with 40 μM biotin. Untransduced JKεKO cells (untr) served as a control. Cells were fixed, permeabilized and stained with an AF647-labeled anti-HA antibody to detect the Ii-hook and measured by flow cytometry. Histograms for total GFP and HA expression are shown. The bar diagrams depict the mean fluorescence intensity (MFI) of total GFP and HA. Significance was analyzed by ordinary one-way ANOVA (n=3; one representative experiment in triplicates is shown).

**(C)** JK wild type (WT) cells were lentivirally transduced with the Ii-εRUSH construct and cultured without biotin (0 min) or in in biotin-containing RPMI medium. Cells were stained with goat anti-GFP and AF680-labeled anti-goat IgG secondary antibodies and TCR surface levels were measured by flow cytometry (n=2).

**(D)** JKεKO Ii-εRUSH cells were cultured without biotin and treated with different concentrations of biotin for 60 min. Cells were stained with an AF647-labeled anti-CD3ε antibody and TCR surface levels were measured by flow cytometry. Bar diagrams depict the mean fluorescence intensity (MFI) of the anti-CD3ε stain. Significance was analyzed by ordinary one-way ANOVA (n=2; one representative experiment in triplicates is shown).

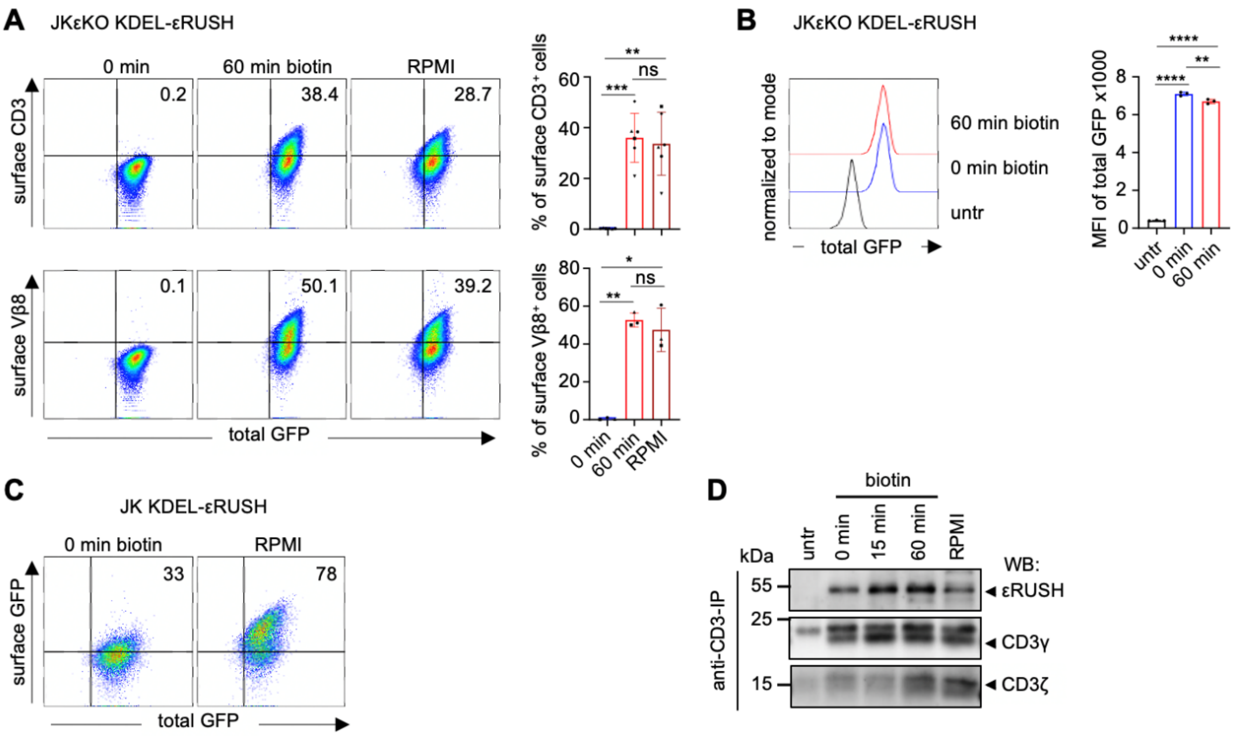


**Figure S2**. **Analysis of the KDEL-εRUSH system in JKεKO and JK WT cells**.

**(A)** JKεKO KDEL-εRUSH cells were cultured without biotin and left untreated (0 min) or treated for 60 min with 40 μM biotin or cultured in RPMI. Cells were stained with an AF647-labeled anti-CD3ε (top panels) and PE-labeled anti-Vβ8 antibody (lower panels). TCR surface levels were measured by flow cytometry. Percentages of surface CD3⁺ and surface TCRβ⁺ within the GFP⁺ cells are plotted. Significance was analyzed by ordinary one-way ANOVA (n=3).

**(B)** JKεKO KDEL-εRUSH cells were cultured without biotin and left untreated (0 min) or treated for 60 min with 40 μM biotin. Untransduced JKεKO cells (untr) served as a control. Cells were measured by flow cytometry. A histogram for GFP expression is shown. The bar diagram depicts the MFI of GFP. Significance was analyzed by ordinary one-way ANOVA (n>3).

**(C)** JK cells were lentivirally transduced with the KDEL-εRUSH construct and cultured without biotin (0 min) or in biotin-containing RPMI medium. Cells were stained with goat anti-GFP and AF680-labeled anti-goat IgG secondary antibodies and TCR surface levels were measured by flow cytometry (n=3).

**(D)** JKεKO KDEL-εRUSH cells were cultured in medium without biotin and left untreated (0 min) or treated for different times with 40 μM biotin. Untransduced JKεKO cells (untr) serve as a control. Cells were lysed and proteins purified by an anti-CD3ε IP. Purified proteins were separated by reducing SDS-PAGE. WB was done using antibodies directed against GFP (εRUSH), CD3γ and CD3ζ (n=3).

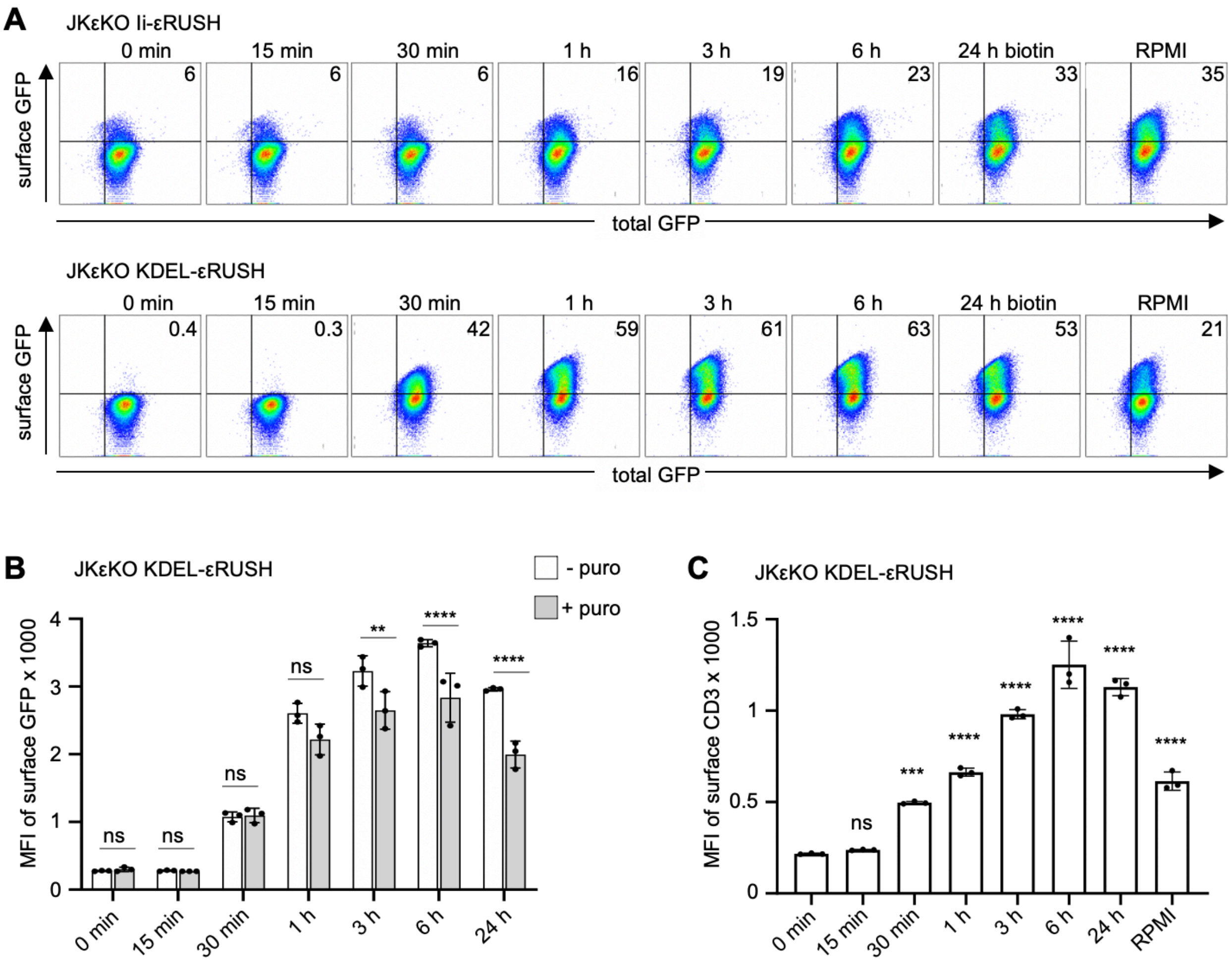


**Figure S3. Kinetics of TCR surface expression in the Ii-εRUSH and KDEL-εRUSH systems.**

**(A)** JKεKO Ii-εRUSH (top panels) and JKεKO KDEL-εRUSH cells (lower panels) were cultured in medium without biotin and left untreated (0 min), treated for different times with 40 μM biotin or were cultured in biotin-containing RPMI medium (RPMI). Cells were stained with AF647-labeled anti-GFP and TCR surface levels were measured by flow cytometry. Dot plots are shown (n=3 for Ii-εRUSH, n>3 for KDEL-εRUSH).

**(B)** JKεKO KDEL-εRUSH cells were cultured and treated as in (A). Before addition of biotin cells were treated for 30 min with puromycin (+puro) and puromycin was kept in culture or cells were left untreated (-puro). Cells were stained with AF647-labeled anti-GFP antibodies and TCR surface levels were measured by flow cytometry. MFI of surface GFP is shown. Significance was analyzed by two-way ANOVA (n=3; one representative experiment in triplicates is shown).

**(C)** JKεKO KDEL-εRUSH cells were cultured and treated as in (A), and stained with an AF647-labeled anti-CD3ε antibody. TCR surface levels were measured by flow cytometry. Bar diagram of the MFI of surface CD3 is shown. Significance to the 0 min sample was analyzed by ordinary one-way ANOVA (n>3; one representative experiment in triplicates is shown).

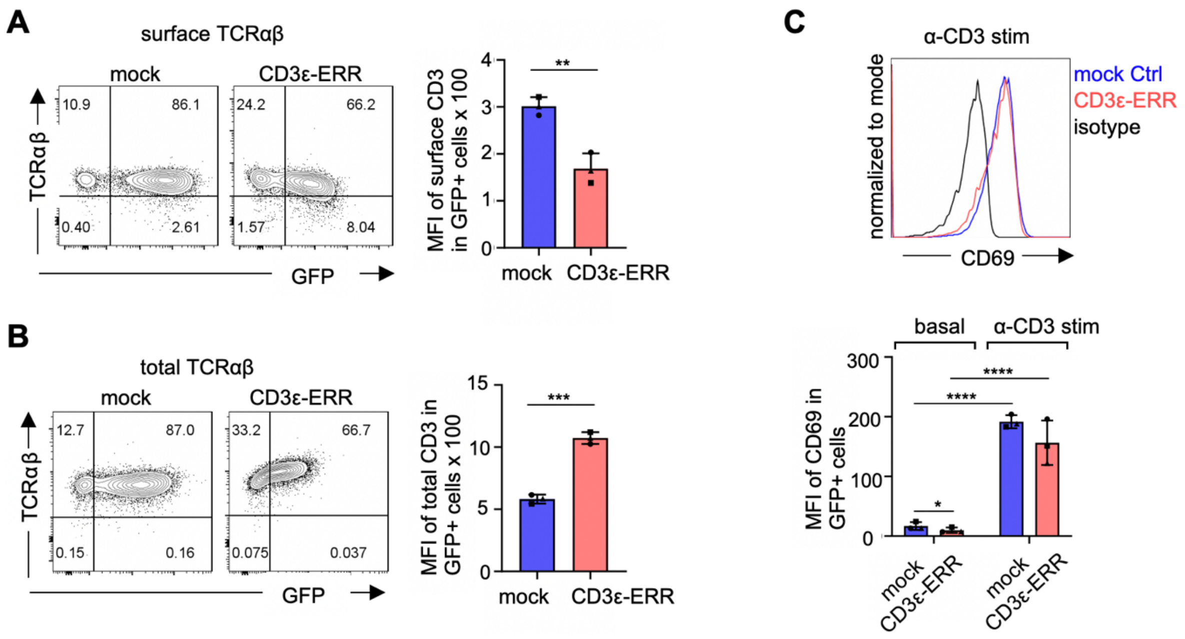


**Figure S4. Analysis of primary human T cells expressing CD3ε-ERR**

**(A)** Cells from figure 6C were stained with an PerCP/Cyanine5.5-labeled anti-TCRαβ antibody and TCR surface levels were measured by flow cytometry. The MFI of TCRαβ in the GFP+ cells is shown.

**(B)** Cells as in (A) were fixed, permeabilized and stained with an PerCP/Cyanine5.5-labeled anti-TCRαβ antibody and measured by flow cytometry. The MFI of TCRαβ in the GFP+ cells is shown.

**(C)** Cells as in (A) were stimulated with a plate-bound anti-CD3ε antibody for 18 hours (α-CD3 stim) or not (basal) and then stained with an PE-labeled anti-CD69 antibody and measured by flow cytometry. Cells stained with an isotype antibody serve as control. The MFI of CD69 in the GFP+ cells is shown.

(A, B and C) Significance was analyzed by paired t-test (n=3 different donors).

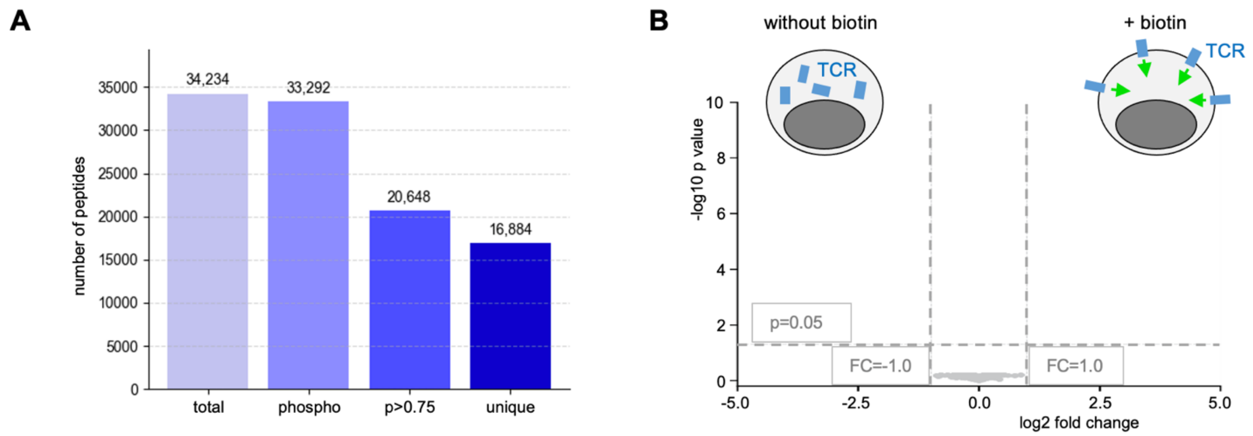


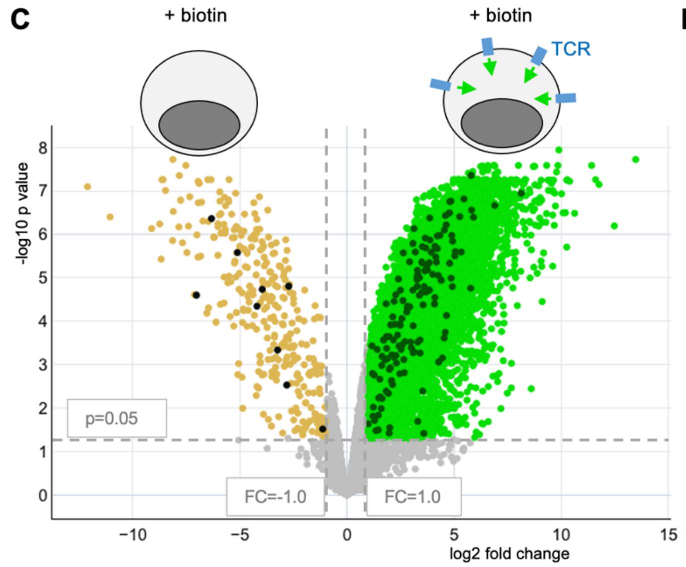


**D**

Phospho-Y sites from (C) that were also detected in figure 7B.
These are part of the basal signal signature.

| protein | phospho-site | log2 fold change |
|---|---|---|
| **CD3δ** | **Y149** | **5.0** |
| **CD3ε** | **Y199** | **4.8** |
| RBMS2 | Y99 | 5.3 |
| ERK2 | Y187 | 4.2 |
| PLCγ1 | Y775 | 4.2 |
| ERK1 | Y204 | 3.8 |
| RIOK2 | Y445 | 3.7 |
| ETF1 | Y96 T99 | 3.5 |
| **CD3ζ** | **Y123** | **2.6** |
| ABI1 | Y213 | 3.4 |
| ZAP70 | Y492 | 3.5 |
| p38α | Y182 | 2.1 |
| **CD3ζ** | **Y142** | **4.3** |
| ZAP70 | Y292 | 4.5 |
| CD5 | Y453 | 4.0 |
| PLCγ1 | Y783 | 4.3 |
| **CD3ζ** | **Y72** | **4.8** |
| PTPN18 | Y389 | 1.2 |
| MAX | Y123 | 2.0 |
| LAT | Y220 | 3.7 |
| LAX1 | Y294 | 1.8 |
| **CD3ζ** | **Y111** | **3.1** |
| c-CBL | Y674 | 1.5 |

**Figure S5. Analysis of the phosphoproteomics data on the basal TCR signal**
**(A)** Mean values of peptide identifications, reporting total number of peptides, phosphopeptides, phosphopeptides with localization probability better than 0.75 and unique phosphopeptides. **(B)** JKεKO KDEL-εRUSH cells were cultured for one hour with (+ biotin) or without biotin. Protein-level intensities recorded before phosphopeptide enrichment in the +biotin condition were compared with those without biotin and displayed as volcano plots based on moderated t-tests. None of the proteins were found significantly changed at the thresholds of <0.05 for p-values and >+1.0 or <-1.0 for the $\log_2$ of the fold change as indicated by the horizonal and vertical dashed lines, respectively. **(C)** JKεKO KDEL-εRUSH and JKεKO cells were each cultured for one hour with (+ biotin). Phosphoproteomics and display as in figure 7A comparing the +/- TCR conditions. Phosphorylations enriched in the cells with a TCR are shown in green and the ones enriched in the cells lacking any TCR are shown in brown. The black dots represent Y phosphorylations. **(D)** Phospho-Y peptides enriched in the JKεKO KDEL-εRUSH sample (with TCR) from (C) and also present in the comparison of JKεKO KDEL-εRUSH cells with and without biotin are shown. These are the phospho-Y sites of the basal signal signature (Table S4).

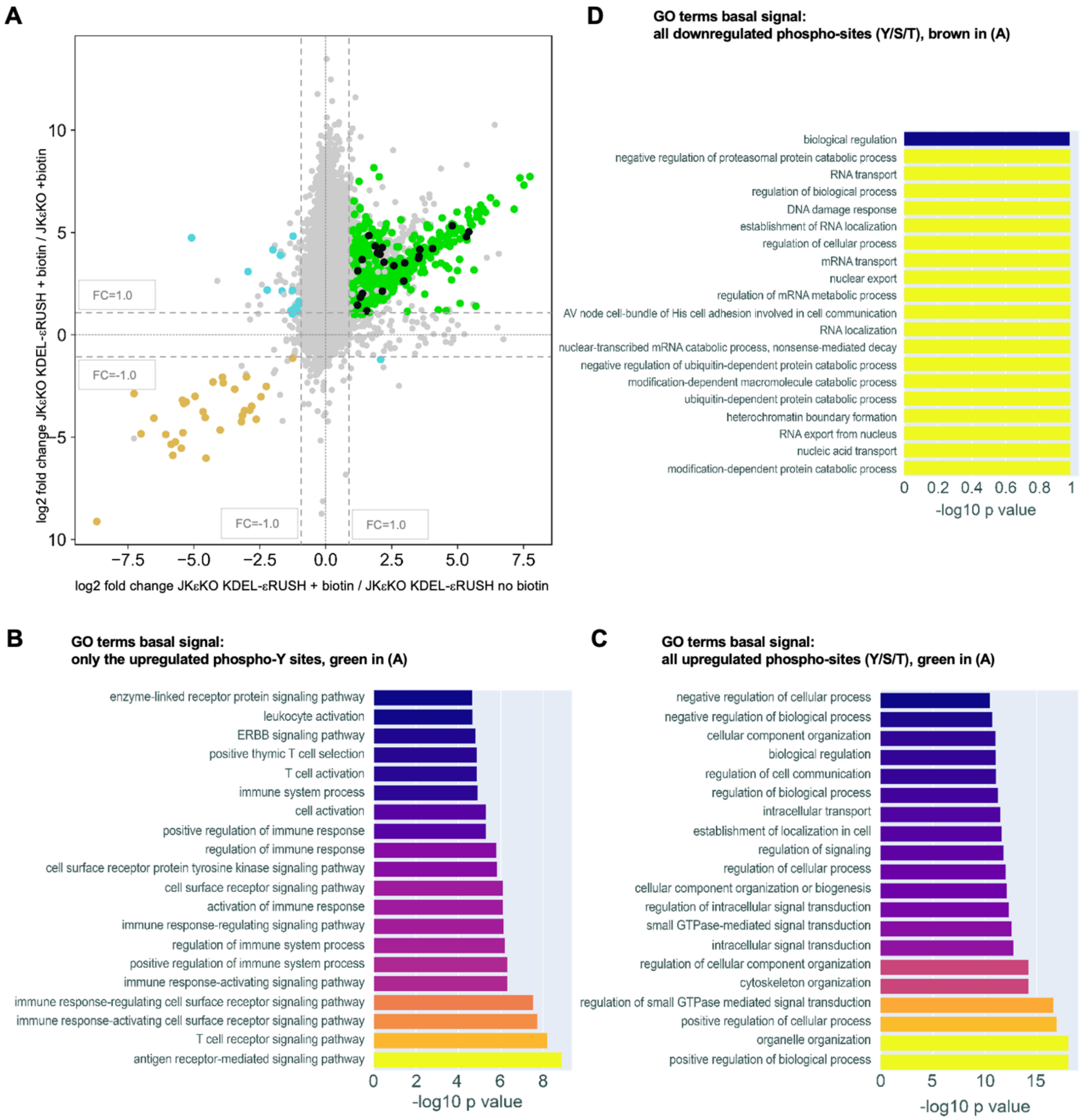


**Figure S6. Identification of the basal signal signature**

**(A)** Fold changes of the basal TCR signal from the +/- biotin comparison as in figure 7A (phosphopeptides from JKεKO KDEL-εRUSH cells with biotin for 1 h versus without biotin, X axis) and of the signal from the +/- TCR comparison in figure S5C (phosphopeptides from JKεKO KDEL-εRUSH cells with biotin versus JKεKO with biotin, Y axis). Phosphopeptides significantly enriched in both signals, with p-values <0.05 and log2 fold changes >1.0, are shown in green, those depleted in both signals are shown in brown. Black dots represent significantly changed tyrosine phosphorylations. Together, the green, brown and black sites identify the basal signal signature. Phosphopeptides with opposite changes in the +/- biotin and the +/- TCR comparison are shown in cyan and all others in grey. **(B)** Gene Ontology (GO) enrichment analysis for phospho-Y upregulated in the basal signal signature in (A). **(C, D)** GO terms associated with all phosphosites upregulated **(C)** or downregulated **(D)** in the basal signal signature.

| protein | phospho-site | log2 fold change | by TCR ligand | activ-ation | comment about the phospho-site | induced by basal signal | enhanced by activation signal |
|---|---|---|---|---|---|---|---|
| WDR48 | Y338 | 6.0 | - | - | unknown | | |
| PP1α | Y306 | 5.1 | - | - | unknown | | |
| RHOGDI2 | Y146 | 4.1 | - | - | unknown | | |
| ERK2 | Y187 | 3.2 | Yes | Yes | located in activation loop | | |
| ERK1 | Y204 | 3.1 | Yes | Yes | located in activation loop | | |
| **CD3δ** | **Y160** | **2.6** | **Yes** | **Yes** | **2nd Y of the ITAM** | unclear | Yes |
| GADS | Y45 | 2.4 | Yes | Yes | relieves inhibition | | |
| **CD3ζ** | **Y83** | **1.9** | **Yes** | **Yes** | **2nd Y of ITAM1** | Yes | Yes |
| RASGRP1 | Y489 | 1.6 | - | - | unknown | | |
| **CD3γ** | **Y171** | **1.5** | **Yes** | **Yes** | **2nd Y of the ITAM** | unclear | Yes |
| LAT | Y255 | 1.4 | Yes | Yes | GRB2 bdg site | | |
| **CD3ε** | **Y188** | **1.3** | **Yes** | **Yes** | **1st Y of the ITAM** | Yes | Yes |
| **CD3ζ** | **Y142** | **1.3** | **Yes** | **Yes** | **1st Y of ITAM3** | Yes | Yes |
| **CD3ζ** | **Y123** | **1.3** | **Yes** | **Yes** | **2nd Y of ITAM2** | Yes | Yes |
| **CD3ε** | **Y199** | **1.1** | **Yes** | **Yes** | **2nd Y of the ITAM** | Yes | Yes |
| LAT | Y220 | 1.0 | Yes | Yes | GRB2 & GADS bdg site | | |
| p38α | Y182 | 1.0 | Yes | Yes | located in activation loop | | |
| ARHGEF6 | Y188 | 1.0 | - | - | unknown | | |
| **CD3ζ** | **Y111** | **0.8** | **Yes** | **Yes** | **1st Y of the ITAM2** | Yes | No |
| **CD3ζ** | **Y153** | **0.6** | **Yes** | **Yes** | **2nd Y of ITAM3** | No | No |
| **CD3ζ** | **Y72** | **0.7** | **Yes** | **Yes** | **1st Y of the ITAM1** | Yes | No |
| **CD3δ** | **Y149** | **0.3** | **Yes** | **Yes** | **1st Y of the ITAM** | Yes | No |
| **CD3ζ** | **S124** | **1.9** | **-** | **-** | unknown | unclear | Yes |

**Figure S7. Analysis of the activation TCR signal**
Selected Y-phosphosites induced by anti-TCR stimulation from figure 7C are shown. Sites known to be induced by ligand binding to the TCR are indicated in the “by TCR ligand” column and the ones known to have an activatory role in ligand-induced TCR signaling in the “activation“ column. CD3 phosphorylation sites induced by basal signaling are marked with “Yes” in the “induced by basal signal” column, and those sites that are then further phosphorylated upon antibody stimulation are marked with “Yes” in the “enhanced by activation signal” column, i.e. those that are more phosphorylated in the activation signal compared to the basal signal.

### Table S1: List of Plasmids

| *plasmid* | *expressed Protein* | *cloning strategy* |
|---|---|---|
| *#65288* | Ii-Str-HA_SBP-EGFP-Ecadherin | From Addgene (Boncompain et al. 2012) |
| *#65278* | Str-KDEL_TNF-SBP-EGFP | From Addgene (Boncompain et al. 2012) |
| *pLVX* | mmp14_zsGreen1 | (Raute et al. 2023) |
| *pOSY-021* | human CD3ε | Sascha Yousefi |
| *pOSY-082* | moxGFP-PIFS-TCRβ | (Yousefi et al. 2019) |
| *p510* | anti-CD19 scFv-CD3γ_GFP | (Baeuerle et al. 2019) |
| *pCJ-01* | Ii-Str-HA_zsGreen1 | Gibson of one fragment: Backbone: pLVX; fragment: PCR with P09fw and P10rv on #65288 |
| *pCJ-02* | Ii-Str-HA_SBP-GFP-CD3ε | Gibson of 3 fragments: Backbone: pCJ-01; fragment-1: PCR with P01fw and P02rv on pOSY-021; fragment-2: PCR with P05fw and P06.1rv on pOSY-082; fragment-3: PCR with P07.1fw and P08rv on pOSY-021 |
| *pCJ-035* | Str-KDEL_SBP-GFP-CD3ε | Gibson of one fragment: Backbone: pCJ-02; fragment: PCR with P70fw and P71rv on #65278 |

### Table S2: List of oligonucleotides

| *primer* | *sequence* |
|---|---|
| *P01fw* | AAAAAACGTCTAGGCCCCCCGAACCACGGGGACGTGGTTTTCCTTTGAAAAACACGATGATAATCCCGGG**ATGCAGTCGGGCACTCACT** |
| *P02rv* | GCACGAAGTTGTTCAAGTTCCCCAGCCAGTCCTTCAACAACGTGTCCACCTCGCCAACCAGTGGTCTTCTCGTCCAT**TTGCCCCCAAACGCCAACT** |
| *P05fw* | ACTGGCTGGGGAACTTGAACAACTTCGTGCACGACTGGAGCATCACCCACAAGGTCAACGTGAACCAGGCAGCGGC**TCCAAGGGCGAGGAGCTGTT** |
| *P06.1rv* | CATTACCATCGCCGGATCC**CTTGTACAGCTCGTCCATGCCGTG** |
| *P07.1fw* | AGCTGTACAAGGGATCCGGC**GATGGTAATGAAGAAATGGGTGG** |
| *P08rv* | ATTTTGTAATCCAGAGGTTGATTGTTCCAGACGCGTGAATTCTTA**GATGCGTCTCTGATTCAGG** |
| *P09fw* | TTGCCGCCAGAACACAGGTGTCGTGACGCGACTAGTGCCACC**ATGCACAGGAGGAGAGCC** |
| *P10rv* | ACGTTAGGGGGGGGGGAGGGAGAGGGGCGGGATCGCGGCCGC**TCACATGGGGACTGGGC** |
| *P70fw* | TGCCGCCAGAACACAGGTGTCGTGACGCGACTAGTGCCACC**ATGGATGTATGCGTCCGTCTTG** |
| *P71rv* | GGGGGGAGGGAGAGGGGCGGGATC**GCGGCCGCTCACAGTTC** |

| *primers for RT-qPCR* | *sequence* |
|---|---|
| *TRAC fw* | CCAGCCCAGAAAGTTCCTGT |
| *TRAC rv* | TAAACCCGGCCACTTTCAGG |
| *B2M fw* | TACACTGAATTCACCCCCAC |
| *B2M rv* | CATCCAATCCAAATGCGGCA |
