## Supplement: Mathematical Modeling for "An ER Retrieval-and-Release System Reveals Ligand-Independent Basal Signaling by Cell Surface TCRs"

**References** **10**

### 1 Overview

This supplement describes the mathematical model of T cell receptor (TCR) production, export, internalization, recycling and degradation. In the experiments (main text), TCR is retained in the endoplasmic reticulum (ER) until it is released at $t = 0$ and moves to the cell surface.

The question is whether surface TCR regulates its own supply and turnover. The proposed model contains two feedbacks by surface TCR: it inhibits ER export and drives recycling. Sections 2–5 describe the data, the model and the estimation, Section 6 the results. All computations use dMod2 0.11.4 [1] with the C++ backend cppDE 0.12.1 [2]. The model cartoon is shown in Figure 4A of the main text.

### 2 Data Preprocessing

Flow cytometry data were obtained from 7 independent release experiments (OPEN), comprising 501 samples. TCR abundance was assessed using a green fluorescent protein (GFP) reporter and, in 1 experiment, additionally by anti-CD3 antibody staining. Each experiment included TCR-negative control cells (CTRL). One experiment that used a different sampling protocol was excluded from the analysis.

Each experiment and readout has its own scale and an additive background. Technical replicates are averaged on the $\log_{10}$ scale (141 experiment means) and aligned with blotIt [3], separately for surface and total TCR:

$$\log_{10} z_k = \log_{10}(\delta_k\, 10^{\eta_{t_k}+s_{g_k}} + 10^{o_{g_k}}) + \epsilon_k, \qquad \epsilon_k \sim \mathcal{N}(0, \sigma^2_{r_k}), \tag{2.1}$$

with the experiment mean $z_k$ at time $t_k$ in group $g_k$ (experiment and readout, 15 groups), the common signal $\eta_t$, scaling $s_g$ and background $o_g$ of group $g$, $\delta_k = 0$ for CTRL and 1 otherwise, and the variability $\sigma_r$ of readout $r$ (surface TCR 0.070 for GFP and 0.066 for anti-CD3, total TCR 0.016 and 0.028, $\log_{10}$ scale).

The aligned data are given on the scale of one reference GFP experiment,

$$y_t = \log_{10}(10^{\eta_t+s_{\text{ref}}} + 10^{o_{\text{ref}}}), \qquad y_{\text{CTRL}} = o_{\text{ref}}, \tag{2.2}$$

the form of the observation function, Eq. (3.2). Standard errors follow by the delta method. This gives 30 data points of surface and total TCR.

ER and post-Golgi TCR were measured by Western blot in 3 blots at 0, 15 and 60 minutes after release, with CTRL cells on every blot, and aligned in the same way (variability 0.044 for ER and 0.036 for post-Golgi TCR). In total the calibration data comprise 38 points (30 flow cytometry, 8 Western blot; Figure S1). 4% of the experiment means deviate by more than two standard deviations; the largest deviation is surface TCR at the last measurement, lower in the second experimental series than in the first (3.31 against 3.54).

### 3 Model

#### 3.1 Compartmental flux model

The model describes the amount of TCR in three compartments: the endoplasmic reticulum (ER), the plasma membrane and the endosome, with the states $\text{TCR}_{\text{ER}}$, $\text{TCR}_{\text{S}}$ and $\text{TCR}_{\text{E}}$ (Figure 4A of the main text). Internalized TCR is either recycled from the endosome to the surface or degraded after transport to lysosomes; the lysosome is not a separate compartment of the model. Each state obeys a mass balance,

$$\frac{\mathrm{d}}{\mathrm{d}t}\text{TCR}_i = \sum_{\text{fluxes into } i} v_k - \sum_{\text{fluxes out of } i} v_k, \tag{3.1}$$

so the model is specified by its fluxes $v_k$ (Table S1).

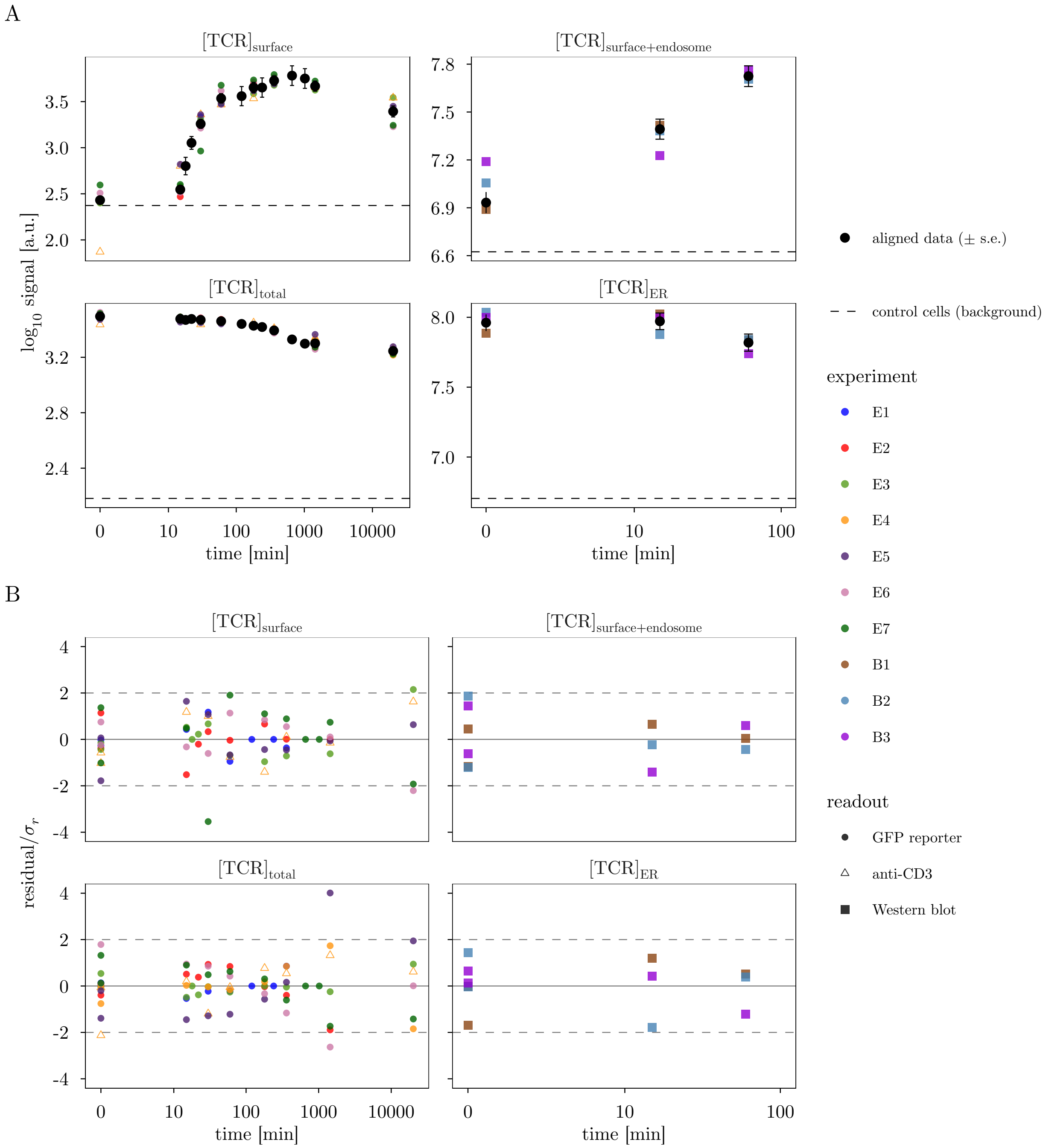

Figure S1: Data: flow cytometry (left, experiments E1–E7) and Western blots (right, B1–B3). (A) Aligned data $y_t$ (black, mean $\pm$ standard error), background $y_{\mathrm{CTRL}}$ (dashed) and the experiment means on the reference scale, colored by experiment. (B) Residuals of the alignment in units of $\sigma_r$; dashed: $\pm 2\sigma_r$. The last measurement is drawn at 14 days.

| Flux | Source | Target | Rate |
|---|---|---|---|
| production | $\emptyset$ | $\mathrm{TCR}_{\mathrm{ER}}$ | $k_P$ |
| ER degradation | $\mathrm{TCR}_{\mathrm{ER}}$ | $\emptyset$ | $k_B\,\mathrm{TCR}_{\mathrm{ER}}$ |
| ER export | $\mathrm{TCR}_{\mathrm{ER}}$ | $\mathrm{TCR}_{\mathrm{S}}$ | $\dfrac{k_E\,\mathrm{TCR}_{\mathrm{ER}}}{1+k^{-}_{\mathrm{S,exp}}\,\mathrm{TCR}_{\mathrm{S}}}$ |
| internalization | $\mathrm{TCR}_{\mathrm{S}}$ | $\mathrm{TCR}_{\mathrm{E}}$ | $\dfrac{k_I\,\mathrm{TCR}_{\mathrm{S}}}{K_m+\mathrm{TCR}_{\mathrm{S}}}$ |
| recycling | $\mathrm{TCR}_{\mathrm{E}}$ | $\mathrm{TCR}_{\mathrm{S}}$ | $k^{+}_{\mathrm{S,rec}}\,\mathrm{TCR}_{\mathrm{S}}\,\mathrm{TCR}_{\mathrm{E}}$ |
| lysosomal degradation | $\mathrm{TCR}_{\mathrm{E}}$ | $\emptyset$ | $\dfrac{k_D\,\mathrm{TCR}_{\mathrm{E}}}{K_m+\mathrm{TCR}_{\mathrm{E}}}$ |

Table S1: Fluxes of the proposed model, $K_m = 10^{-4}$.

**ER export** follows first-order kinetics ($k_E$). Exported TCR reaches the surface directly: a Golgi chain of one, three or five compartments gives the same fit (data terms -149.22, -149.22, -149.22 in an earlier model version calibrated to flow cytometry alone).

**Internalization and lysosomal degradation** are saturated with $K_m = 10^{-4}$, far below the amounts in the experiment, so they run at their capacities $k_I$ and $k_D$ (zeroth order). **ER degradation** is first order ($k_B$, fixed by the balance before release, Section 4).

**Regulation.** An inhibition divides a rate by $1+k^{-}\,\mathrm{TCR}$, a promotion multiplies it by $1+k^{+}\,\mathrm{TCR}$. Feedbacks are named by effect ($\pm$), regulating TCR (S: surface) and flux. The proposed model has two: surface TCR inhibits ER export ($k^{-}_{\mathrm{S,exp}}$) and promotes recycling ($k^{+}_{\mathrm{S,rec}}$). Recycling has no basal rate; a basal rate and the promotion were not separately identifiable.

#### 3.2 Observation function

Flow cytometry measures surface and total TCR, the Western blots TCR in the ER form and in the post-Golgi form (TCR that has left the ER). All four are analyzed on a $\log_{10}$ scale:

$$\begin{aligned} y_{\text{surface}} &= \log_{10}(\mathrm{TCR}_{\mathrm{S}} + a_{\text{surface}}) + b_{\text{surface}}, \\ y_{\text{total}} &= \log_{10}(\mathrm{TCR}_{\mathrm{ER}} + \mathrm{TCR}_{\mathrm{S}} + \mathrm{TCR}_{\mathrm{E}} + a_{\text{total}}) + b_{\text{total}}, \\ y_{\text{ER}} &= \log_{10}(\mathrm{TCR}_{\mathrm{ER}} + a_{\text{ER}}) + b_{\text{ER}}, \\ y_{\text{post-Golgi}} &= \log_{10}(\mathrm{TCR}_{\mathrm{S}} + \mathrm{TCR}_{\mathrm{E}} + a_{\text{post-Golgi}}) + b_{\text{post-Golgi}}. \end{aligned} \tag{3.2}$$

The backgrounds $a$ (autofluorescence, unspecific staining, blot background) are determined by the cells without TCR; the offsets $b$ are scalings on the linear scale.

#### 3.3 Structural identifiability and gauge fixing

All measurements are in arbitrary units. Apart from the fixed $K_m$, the model is invariant under the scaling, for $c > 0$,

$$\begin{aligned} x(t) &\mapsto c\,x(t), & \mathrm{TCR}_{\mathrm{ER}}(0) &\mapsto c\,\mathrm{TCR}_{\mathrm{ER}}(0), & k_P,\ k_I,\ k_D &\mapsto c\,k_P,\ c\,k_I,\ c\,k_D, \\ a_y &\mapsto c\,a_y, & b_y &\mapsto b_y - \log_{10} c, & k^{-}_{\mathrm{S,exp}},\ k^{+}_{\mathrm{S,rec}} &\mapsto k^{-}_{\mathrm{S,exp}}/c,\ k^{+}_{\mathrm{S,rec}}/c, \end{aligned} \tag{3.3}$$

where $x$ are all TCR amounts and $a_y$, $b_y$ the background and scaling of each observable; inserting Eq. (3.3) into Table S1 and Eq. (3.2) leaves all observables unchanged. We fix the gauge $b_{\text{surface}} = 0$, which defines the

| Inner parameter | OPEN | CTRL |
|---|---|---|
| $k_P$ | $10^{\theta_{k_P}}$ | 0 |
| $k_B$ | $k_P/\mathrm{TCR}_{\mathrm{ER}}(0)$ | – |
| $k_E,\ k_I,\ k_D,\ k^-_{\mathrm{S,exp}},\ k^+_{\mathrm{S,rec}}$ | $10^{\theta}$ | $10^{\theta}$ |
| $a_{\mathrm{surface}},\ a_{\mathrm{total}},\ a_{\mathrm{ER}},\ a_{\mathrm{post\text{-}Golgi}}$ | $10^{\theta}$ | $10^{\theta}$ |
| $b_{\mathrm{total}},\ b_{\mathrm{ER}},\ b_{\mathrm{post\text{-}Golgi}}$ | $\theta_b$ | $\theta_b$ |
| $b_{\mathrm{surface}}$ | 0 (gauge, Section 3.3) | 0 |
| $\mathrm{TCR}_{\mathrm{ER}}(0)$ | $10^{\theta_{\mathrm{TCR_{ER}}}}$ | 0 |
| other initial values | 0 | 0 |

Table S2: Parameter transformation from the estimated (outer) parameters $\theta$ to the model (inner) parameters, per condition.

arbitrary unit (a.u.) of the TCR amounts.

### 4 Conditions, initial values and parameters

The two conditions of the data (Section 2) enter the model as follows.

- **CTRL** are cells without TCR: all states and $k_P$ are zero, and the observables reduce to $\log_{10} a + b$.
- **OPEN** is the release experiment. Before release, only $\mathrm{TCR}_{\mathrm{ER}}(0)$ is nonzero, in balance between production and ER degradation.

This balance gives

$$k_B = \frac{k_P}{\mathrm{TCR}_{\mathrm{ER}}(0)}. \tag{4.1}$$

The last measurement, at least 14 days after release, is fitted at 14 days. All parameters except $b$ are estimated on a $\log_{10}$ scale (Table S2).

The model has 14 parameters: seven kinetic and seven observation parameters.

### 5 Parameter estimation

#### 5.1 Objective function

With independent Gaussian measurement errors of known standard deviation $\sigma_i$, the negative twofold log-likelihood is

$$\ell_{\mathrm{data}}(\theta) = \sum_{i=1}^{n} \left[ \frac{\left(y_i - g(x(t_i,\theta),\theta)\right)^2}{\sigma_i^2} + \log(2\pi\sigma_i^2) \right], \tag{5.1}$$

with $n = 38$ data points. Since the cells are back at homeostasis at the last measurement, the objective contains a steady-state constraint at $t_{14} = 14$ days,

$$\ell(\theta) = \ell_{\mathrm{data}}(\theta) + \sum_j \left( \frac{f_j(x(t_{14},\theta),\theta)}{s_{\mathrm{s}}} \right)^2, \qquad s_{\mathrm{s}} = 10^{-3}\ \mathrm{min}^{-1}. \tag{5.2}$$

which treats the right-hand sides $f_j$ at 14 days as data zero with standard deviation $s_{\mathrm{s}}$ (a.u./min). There is no prior. Likelihood ratio tests [4] and profiles use $\ell_{\mathrm{data}}$.

##### 5.2 Optimization

The model and its sensitivities are integrated by cppDE [2] (relative and absolute tolerances $10^{-6}$, $10^{-8}$), and $\ell$ is minimized with the trust-region optimizer of dMod2 without bounds, from 200 starts drawn normally (standard deviation 3, $\log_{10}$ scale) around the orders of magnitude of the parameters. 175 fits converged; 9 reach the maximum likelihood estimate (MLE) $\hat{\theta}$ within 0.1. The next local optimum has an objective value higher by 102.5 (Figure S2), so the MLE is found reliably.

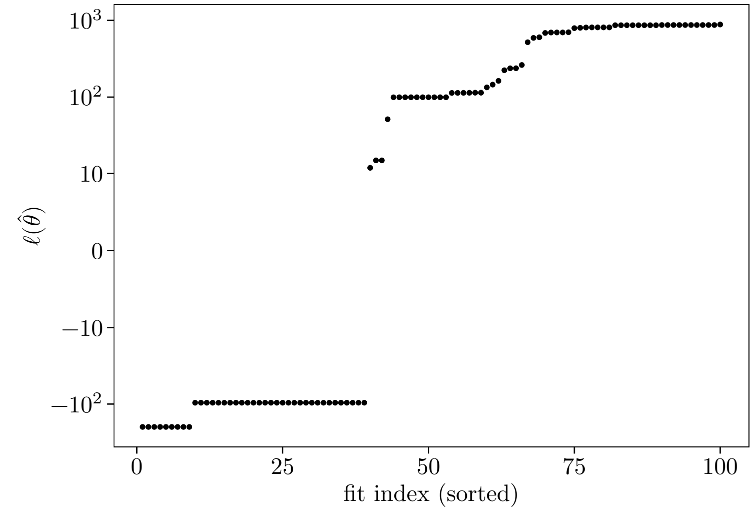

Figure S2: The first 100 converged fits, sorted by their objective value $\ell(\hat{\theta})$, Eq. (5.2).

#### 6 Results

##### 6.1 Fit to the data

The proposed model describes all four observables (Figure S3; $\ell_{\text{data}} = -162.70$).

##### 6.2 All parameters are identifiable

The profile likelihood [5] re-optimizes all other parameters along one parameter and records $\Delta\ell_{\text{data}}$; the 95% confidence interval contains all values with $\Delta\ell_{\text{data}} < 3.84$. All 14 parameters have finite intervals (Figure S4, Table S3), and the steady-state constraint stays at its minimum along all profiles (largest increase 0.00).

**Each feedback is required.** Both feedback profiles are bounded towards zero, so neither feedback can be removed without a significant loss of fit.

##### 6.3 Dynamics of the proposed model

After release, the export flux first stays below the internalization capacity $k_I$: exported TCR is internalized at once and the endosome fills. Recycling grows with the surface and the endosomal amount and eventually exceeds $k_I$, so surface TCR rises sharply about 15 minutes after release. Surface TCR then throttles export and drives recycling; the ER empties slowly and surface TCR overshoots before it settles.

The saturated rates per TCR depend on the amounts. At the surface maximum, $\text{TCR}_\text{S} = 5406$,a.u., internalization runs at $0.018\,\text{min}^{-1}$ per surface TCR, and at homeostasis at $0.037\,\text{min}^{-1}$, close to the $0.012\,\text{min}^{-1}$ measured for the TCR [6]. Recycling, $k^{+}_{\text{S,rec}}\,\text{TCR}_\text{S}$ per endosomal TCR, is $1.23\,\text{min}^{-1}$ at homeostasis, faster than the $0.055\,\text{min}^{-1}$ reported there: in the model the endosome is a small, fast compartment (52,a.u. at homeostasis).

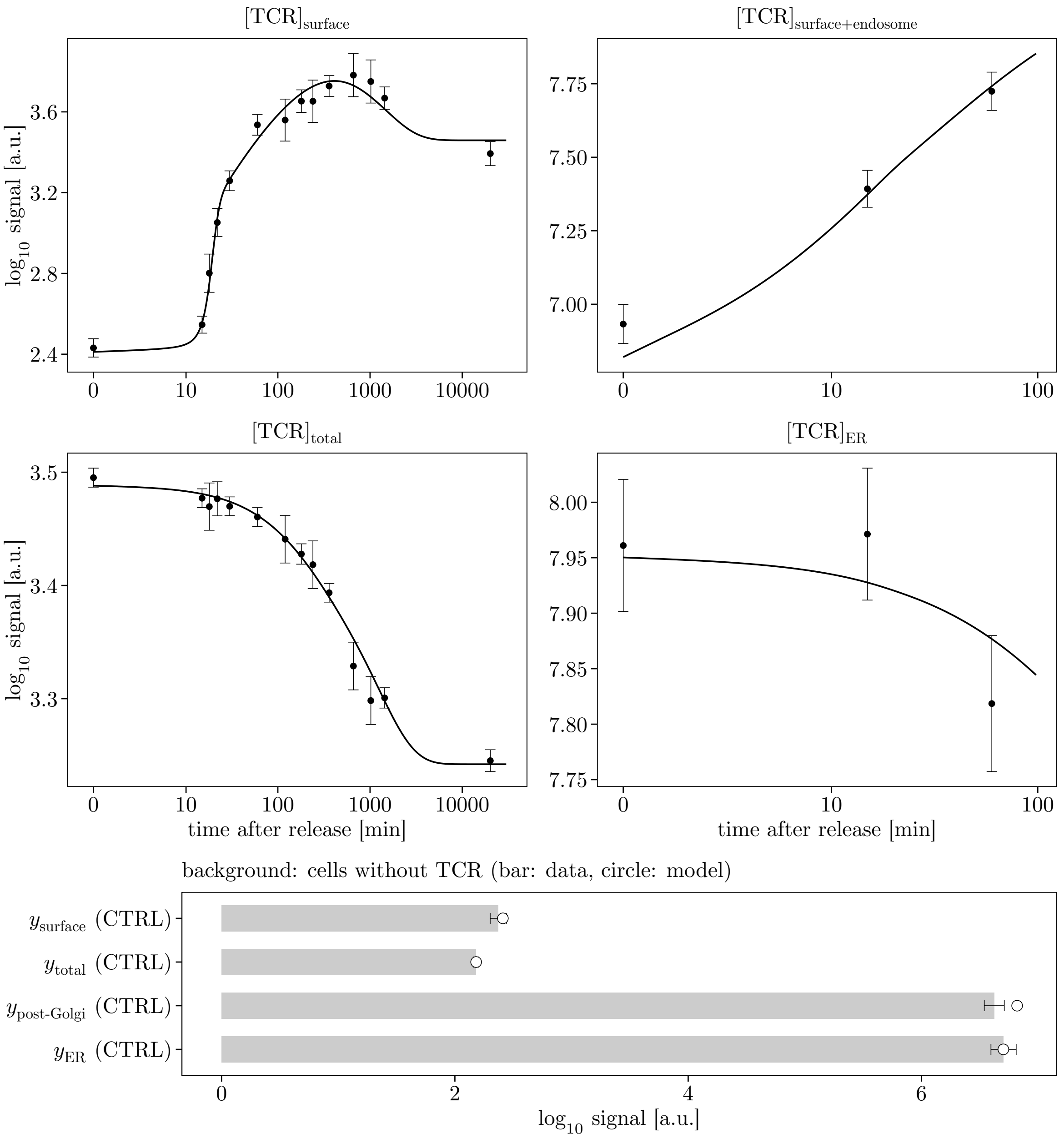

Figure S3: Model at the MLE (line) and data (mean ± standard error). Flow cytometry left, Western blots right. Bottom: background of cells without TCR (bar: data, circle: model). The last measurement is drawn at 14 days.

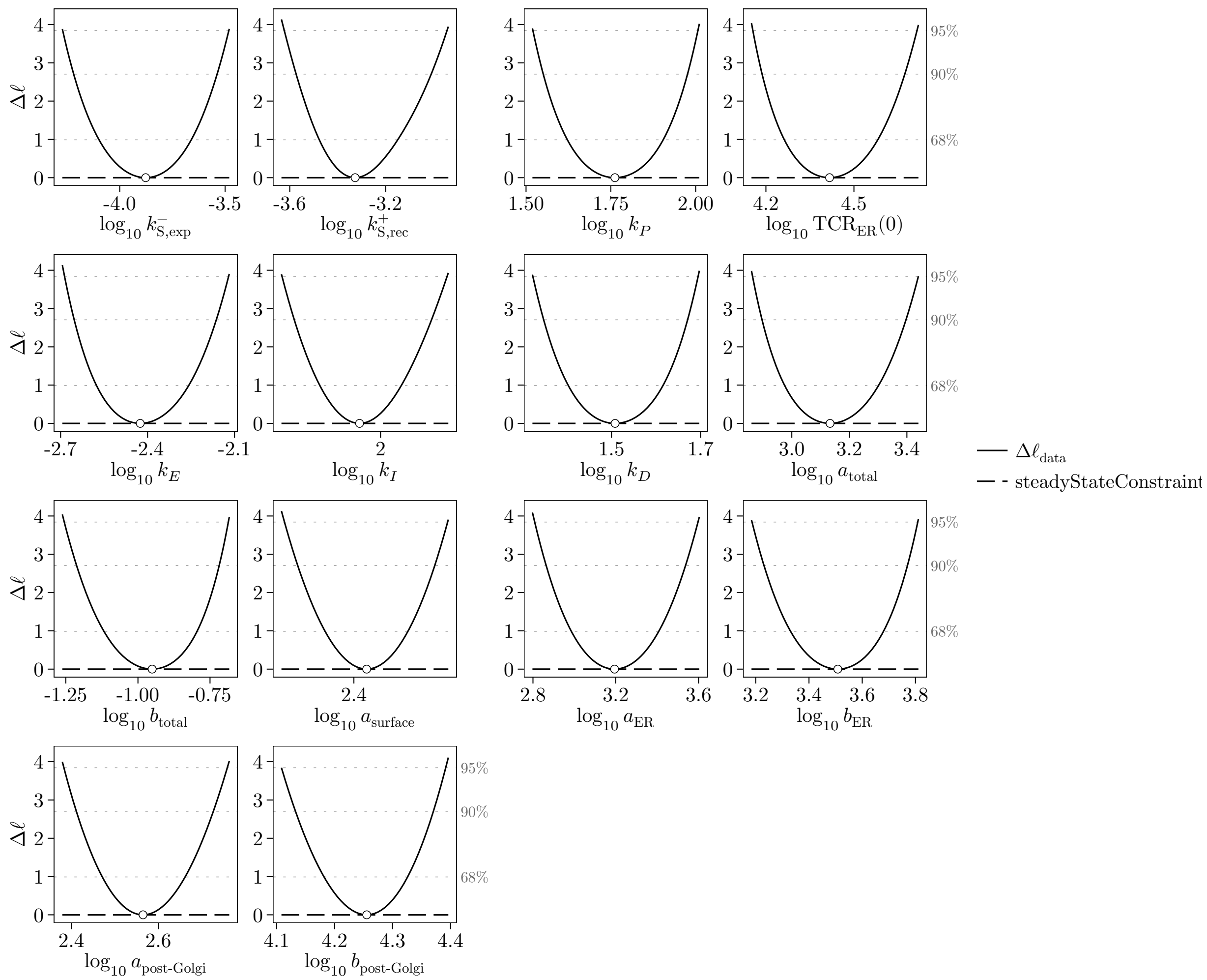

Figure S4: Profile likelihoods: data term (solid) and steady-state constraint (dashed). Dotted: 68%, 90% and 95% thresholds; circle: MLE.

#### 6.4 Tests of the kinetics

Free Michaelis constants for all four fluxes, or additional first-order terms for internalization and degradation, do not improve the fit significantly; purely first-order internalization and degradation fit worse (Table S4).

#### 6.5 No further feedback is required

Each of the other 18 feedbacks on production, export, internalization, recycling and degradation by surface or ER TCR was added on its own, refitted from 52 starts and tested with one degree of freedom, $\Lambda = \ell_{\text{data}}(\text{proposed}) - \ell_{\text{data}}(\text{extended})$. No feedback passes the 95% threshold (Figure 4C of the main text); the largest improvement is $\Lambda = 0.53$ (surface TCR promotes ER export, $p = 0.47$).

| Parameter | unit | $\hat{\theta}$ | lower | upper |
|---|---|---|---|---|
| $k^{-}_{\mathrm{S,exp}}$ | 1/a.u. | -3.88 | -4.27 | -3.48 |
| $k^{+}_{\mathrm{S,rec}}$ | 1/(a.u. min) | -3.33 | -3.63 | -2.94 |
| $k_P$ | a.u./min | 1.76 | 1.52 | 2.01 |
| $\mathrm{TCR}_{\mathrm{ER}}(0)$ | a.u. | 4.42 | 4.15 | 4.72 |
| $k_E$ | 1/min | -2.43 | -2.69 | -2.12 |
| $k_I$ | a.u./min | 1.98 | 1.91 | 2.06 |
| $k_D$ | a.u./min | 1.51 | 1.32 | 1.70 |
| $a_{\mathrm{surface}}$ | a.u. | 2.41 | 2.33 | 2.48 |
| $a_{\mathrm{total}}$ | a.u. | 3.13 | 2.86 | 3.44 |
| $b_{\mathrm{total}}$ | – | -0.95 | -1.26 | -0.68 |
| $a_{\mathrm{ER}}$ | a.u. | 3.19 | 2.80 | 3.60 |
| $b_{\mathrm{ER}}$ | – | 3.51 | 3.18 | 3.81 |
| $a_{\mathrm{post\text{-}Golgi}}$ | a.u. | 2.56 | 2.38 | 2.76 |
| $b_{\mathrm{post\text{-}Golgi}}$ | – | 4.26 | 4.11 | 4.40 |

Table S3: Parameters of the proposed model on the $\log_{10}$ scale ($b$ linear), with the bounds of the 95% confidence intervals from the profile likelihood.

| Kinetics | parameters | $\ell_{\mathrm{data}}$ | $\Lambda$ | df | $p$ |
|---|---|---|---|---|---|
| proposed model (saturated internalization and degradation) | 14 | -162.70 | – | – | – |
| internalization and lysosomal degradation of first order | 14 | -160.16 | – | 0 | – |
| Michaelis constants of export, internalization, recycling and degradation estimated | 18 | -163.36 | 0.66 | 4 | 0.96 |
| lysosomal degradation of zeroth and first order | 15 | -163.57 | 0.87 | 1 | 0.35 |
| internalization and lysosomal degradation of zeroth and first order | 16 | -163.57 | 0.87 | 2 | 0.65 |

Table S4: Kinetic alternatives. $\Lambda = \ell_{\mathrm{data}}(\mathrm{proposed}) - \ell_{\mathrm{data}}(\mathrm{alternative})$ for nested alternatives (df: additional parameters), $p$ from the $\chi^2_{\mathrm{df}}$ distribution. The first-order alternative has the same number of parameters; its data term is higher by 2.55.
